# A common tag nucleotide variant in *MMP7* promoter increases risk for hypertension via enhanced interactions with CREB transcription factor

**DOI:** 10.1101/568774

**Authors:** Lakshmi Subramanian, Sakthisree Maghajothi, Mrityunjay Singh, Kousik Kesh, Kalyani Ananthamohan, Saurabh Sharma, Madhu Khullar, Suma M. Victor, Snehasikta Swarnakar, Shailendra Asthana, Ajit S. Mullasari, Nitish R. Mahapatra

## Abstract

MMP7 (Matrilysin), a potent extracellular matrix degrading enzyme with wide substrate specificity, is emerging as a new regulator of cardiovascular diseases including coronary artery disease and atherosclerosis. However, potential contributions of *MMP7* genetic variations to hypertension remain unknown. In this study, we first probed for the association of a tag single nucleotide polymorphism (SNP) in the *MMP7* gene promoter (-181A/G; rs11568818) with hypertension in an urban south Indian population (n=1517). The heterozygous A/G genotype showed a strong association with hypertension as compared to the A/A wild-type genotype (OR=1.641, 95% CI=1.276-2.109; p=1×10^−4^); AG genotype carriers also displayed significantly higher diastolic blood pressure and mean arterial pressure than AA genotype subjects. The study was replicated in a north Indian population (n=977) as well (OR=1.520, 95% CI =1.106-2.090; p=0.01). Transient transfection experiments using *MMP7* promoter-luciferase reporter constructs revealed that the variant -181G allele conferred greater promoter activity than the -181A allele. Computational prediction and structure-based conformational and molecular dynamics simulation studies suggested higher binding affinity for the transcription factor CREB to the -181G promoter. In corroboration, over-expression/down-regulation of CREB and chromatin immunoprecipitation experiments provided convincing evidence for stronger binding of CREB with the -181G promoter. Further, the -181G promoter also displayed an enhanced response to hypoxia and epinephrine-treatment. The higher promoter activity of -181G allele also translated to increased MMP7 protein levels. Indeed, *MMP7*-181A/G heterozygous individuals displayed elevated plasma MMP7 levels which positively correlated with blood pressure. In conclusion, the *MMP7* A-181G promoter SNP increased expression of MMP7 under pathophysiological (such as hypoxic stress and catecholamine excess) conditions via increased interactions with the transcription factor CREB and enhanced the risk for hypertension in its carriers.

## INTRODUCTION

Matrix metalloproteinases (MMPs) are enzymes that degrade the components of extracellular matrix (ECM), basement membrane thereby contributing to tissue remodelling and developmental processes. They are also known to cleave bioactive molecules, chemokines, cytokines, growth factors involved in physiological processes including inflammation, angiogenesis, wound healing etc.^1^ To date, 24 different MMPs have been identified and classified into collagenases, stromelysins, gelatinases, matrilysins and membrane-type MMPs based on their domain and substrate specificities. MMP gene expression is primarily regulated at the transcriptional level and their activities are modulated by zymogen activation and specific inhibitors called tissue inhibitors of matrix metalloproteinases and non-specific inhibitors like α2-macroglobulin.^2^

MMP7, the smallest (molecular weight: 28 kDa) matrix metalloproteinase, is a secreted matrilysin with specificity for a broad range of substrates (viz. fibronectin, elastin, type IV collagen and proteoglycans). MMP7 is also known to cleave pro-heparin binding epidermal growth factor, Fas ligand, E-cadherin, insulin-like growth factor binding protein, β2 adrenergic receptor, VEGFR2, TNF-α and activate other MMPs like pro-MMP2 and pro-MMP9.^3^ Serum/Plasma MMP7 levels are found to be elevated in several types of cancers including gastric cancer, colorectal cancer, ovarian cancer, lung cancer, prostate cancer etc, atherosclerosis, hypertension, chronic kidney disease and diabetes.^4-7^ Since uncontrolled proteolytic processes leading to vascular remodelling acts as an important determinant of cardiovascular complications including hypertension, myocardial infarction and stroke, MMPs are expected and shown to serve as potential mediators in these disease states.^8^ For instance, knock-down of *MMP7* in the spontaneous hypertensive rat model attenuated hypertension.^9^ MMP7 was found to be an upstream transcriptional activator of MMP-2 and knock-down of *MMP7* prevented the progression of angiotensin II (Ang II)-induced hypertension and cardiac hypertrophy.^10^ Genetic polymorphisms in the *MMP7* have been shown to be associated with coronary artery disease (CAD), acute MI, multiple sclerosis, rheumatoid arthritis, and several cancers.^7, 11^ The *MMP7* promoter SNPs A-181G and C-153T, among the most widely studied SNPs, were shown to be functional as they displayed allele-specific effects and modulated gene expression due to differential interaction of nuclear binding proteins.^11,12^ However, potential association of regulatory variants in *MMP7* gene promoter with hypertension has not been reported so far. We hypothesize that association studies in ethnically distinct populations and functional characterization of genetic variants in *MMP7* promoter may help understand the mechanisms behind dysregulated MMP7 levels in cardiovascular diseases (CVDs) as transcriptional regulation plays a key role in controlling *MMP7* gene expression levels.

In this study, we probed for association of the common tag SNP A-181G (rs11568818) in the *MMP7* gene promoter using a case-control approach in two geographically distinct Indian populations. The A-181G polymorphism increased risk for hypertension in both study populations. Functional characterization revealed that the variant *MMP7*-181G promoter displayed higher promoter activity than the wild-type *MMP7*-181A promoter. Systematic computational, cellular and molecular studies established allele-specific regulation by the transcription factor cAMP response element-binding protein (CREB) in enhancing the *MMP7*-181G promoter activity in basal as well as in pathophysiological conditions. In corroboration, the carriers of -181A/G heterozygous genotype displayed elevated plasma MMP7 levels and higher blood pressure justifying the functional role of this polymorphism to enhance the risk for hypertension.

## MATERIALS AND METHODS

The detailed methods are provided in the Supplementary Information.

### Human Subjects

The primary study population comprised of 862 hypertensive and 655 normotensive individuals from an urban Chennai population recruited at the Madras Medical Mission (MMM), Chennai. A replication study was carried out with 519 hypertensive and 458 normotensive individuals from Chandigarh recruited at the Postgraduate Institute of Medical Education and Research (PGIMER), Chandigarh. Each volunteer gave an informed written consent and the study was approved by the institutional ethics Committee at Indian Institute of Technology Madras. Demographic, physiological and biochemical parameters of both the study populations are listed in the Tables S1 and S2.

### Genotyping of *MMP7* -181A/G polymorphism

Genomic DNA was isolated from heparin/EDTA-anticoagulated blood samples using Flexigene DNA kit. The promoter region of *MMP7* comprising of -302 bp to -153 bp (NCBI accession number: NM_002423.4) was PCR-amplified using specific primers, purified and digested with EcoRI to detect the genotypes at the -181 bp position (Fig. S1).

### Estimation of biochemical parameters

Common biochemical parameters such as glucose, lipid profile, urea, creatinine, hemoglobin, sodium and potassium levels in plasma were estimated by standard assays. Blood pressure was measured in the sitting position using a brachial oscillometric cuff by experienced nursing staff and the readings were recorded in triplicate and averaged. Plasma MMP7 levels were measured by an ELISA kit.

### Cloning and mutagenesis

The *MMP7*-181G and *MMP7*-181A constructs were generated in pGL3-Basic vector using specific primers; these constructs harbor -230 bp to +22 bp region of *MMP7* gene. *MMP7*-181A/G promoter-cDNA constructs were generated by replacing the firefly luciferase cDNA in *MMP7*-181A and *MMP7*-181G promoter-reporter constructs with MMP7 cDNA.

### Cell lines, transfection and reporter assays

Human neuroblastoma IMR-32, SH-SY5Y, rat cardiomyoblast H9c2, and mouse neuroblastoma N2a cell lines were obtained from the National Center for Cell Sciences, Pune, India. All transfections were carried out using Targefect F2 transfection reagent. *MMP7* promoter-reporter constructs and β-galactosidase (β-gal) expression plasmid (internal control) were transfected into these cell lines. Luciferase and β-galactosidase assays were performed as reported previously^13^ after 24-30 hrs of transfection and promoter activities were expressed as luciferase/β-galactosidase readings.

Co-transfection experiments with CREB and KCREB expression plasmids^14^, CREB dicer substrate siRNA oligos and treatments (epinephrine and hypoxia) were carried out in IMR-32 and H9c2 cells along/after transfection with *MMP7*-181G and *MMP7*-181A promoter-reporter constructs. Luciferase activity and total protein levels were estimated and promoter activities were expressed as luciferase/μg of protein. Experiments were also carried out to imitate the homozygous and heterozygous conditions *in cella* with *MMP7* promoter-reporter constructs and by transfecting *MMP7* promoter-cDNA constructs to estimate MMP7 levels *in vitro*.

### Western blotting

Immunoblotting experiments were performed to detect over-expression or down-regulation of CREB, phospho-CREB and HIF-1α after transfection experiments/epinephrine treatment/hypoxia and to measure the influence of *MMP7*-181G and -181A promoter variants on the MMP7 levels *in vitro* using specific antibodies.

### Chromatin immunoprecipitation (ChIP) assays

ChIP assays were carried out in N2a and H9c2 cells transfected with *MMP7*-181G and *MMP7*-181A promoter-reporter constructs with/without treatment with epinephrine (5 μM) or hypoxia following our recently reported protocol.^5^ Immunoprecipitation reactions were carried out using CREB/phospho-CREB antibodies with IgG serving as negative control. The region encompassing the *MMP7*-181A/G promoter was amplified by qPCR using specific primers and the amount of DNA immunoprecipitated in case of -181G and -181A alleles due to CREB binding were quantified by the fold enrichment method relative to IgG signal.

### Computational analysis

The homology model of CREB1 was built using MODELLER v11.9^15^, using the template 1DH3 (285-339, *Mus musculus*, mCREB1, resolution: 2.8 Å). The *MMP7*-181G and *MMP7*-181A promoter DNA models were built using 3DDART^16^ and SCFBio (http://www.scfbio-iitd.res.in/research/drugdna.html). Protein–DNA docking was performed with HDOCK^17^ and HADDOCK^18^ tools. Molecular dynamics (MD) simulation study for all CREB1-*MMP7* (protein-DNA) complexes/models were carried out using Desmond^19, 20^ tool in Maestro.

### Data presentation and statistical analysis

Phenotypic measurements of the study population were expressed as mean ± SEM. Genotype-phenotype associations were tested by one-way ANOVA with post-hoc tests or Levene’s test for equality of variances followed by two-tailed t-test, as appropriate, using Statistical Package for Social Sciences (SPSS). Promoter-reporter transfections results were expressed as mean ± SEM from representative experiments. Statistical significance was calculated by Student’s t-test, one-way ANOVA or two-way ANOVA with post-hoc tests, as applicable, using Prism 5 program.

## RESULTS

### Identification and linkage disequilibrium analysis of SNPs in the *MMP7* promoter

SNPs occurring at a frequency ≥ 1% in the 5 kb *MMP7* promoter in the south Asian population of the 1000 Genomes project were identified from dbSNP database (Fig. 1 and Table S3). Of the 10 SNPs identified, 7 were found to be common polymorphisms (occurring at a frequency of ≥ 5%). Pair-wise linkage disequilibrium (LD) analysis was carried out to predict the non-random association of the alleles of these ten SNPs in the *MMP7* promoter and to shortlist tag SNPs, if any. Eight of the ten SNPs constituted a haplotype block suggesting that these alleles could be inherited together (Fig. 1B). Additionally, tag SNP prediction among these SNPs carried out using LD tag SNP selection tool from SNPinfo^21^ web server predicted the rs11568818 (*MMP7* A-181G) polymorphism occurring at a frequency of 0.36 (minor allele frequency (MAF)) in the overall 1000 Genomes population and 0.43 in the south Asian super-population as a tag SNP. The variant at -181 bp was found to be in LD with SNPs at -1378 bp (rs17098318) and -1773 bp (rs17881620) (Fig. 1B).

**Figure 1.**
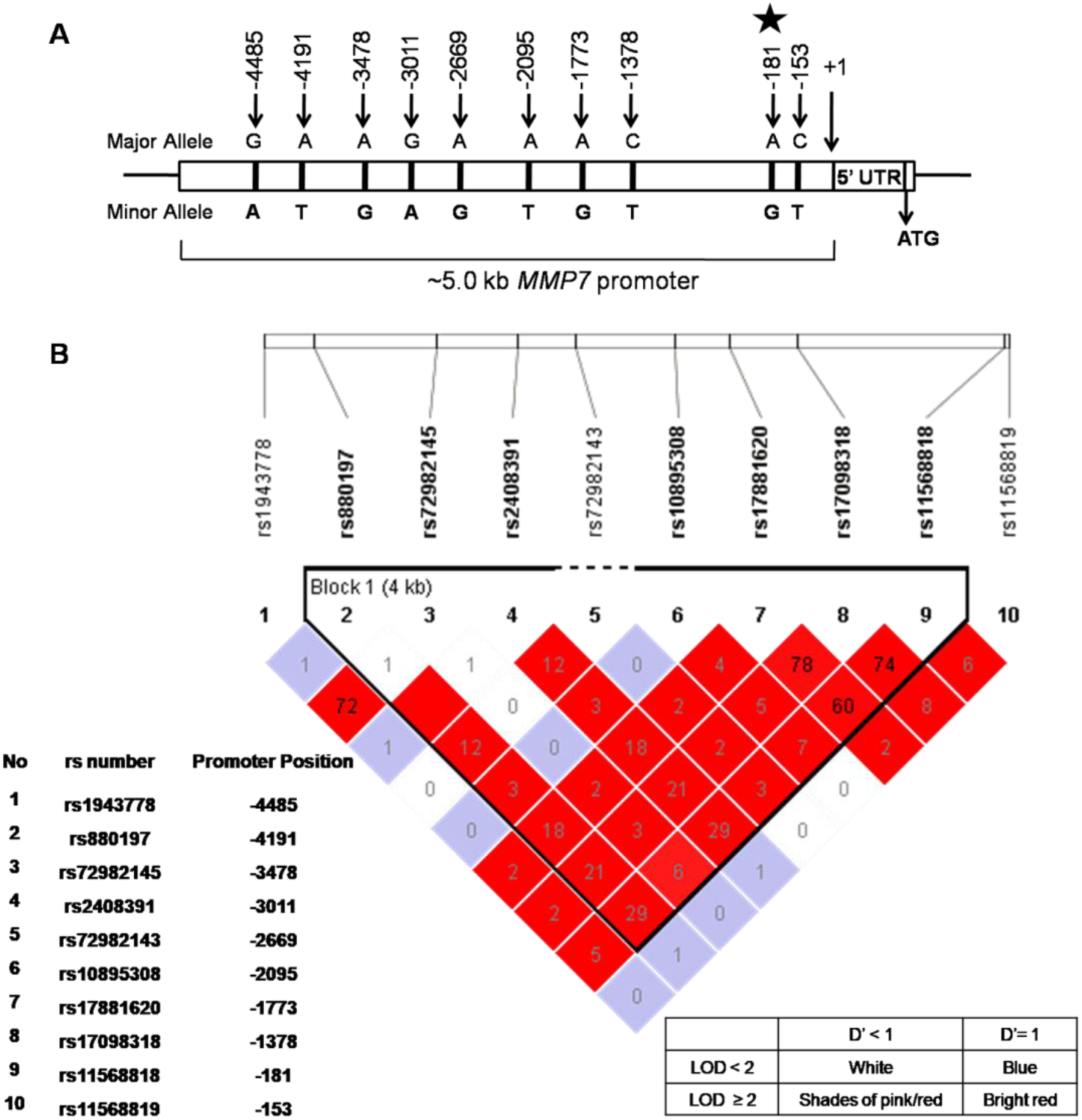
Common promoter polymorphisms in *MMP7*. **(A)** Schematic showing the common polymorphisms in the 5 kb promoter region of *MMP7*. *MMP7* A-181G polymorphism was identified as a tag SNP. **(B)** *MMP7* promoter polymorphisms and linkage disequilibrium (LD). LD plot of common polymorphisms in 5 kb upstream region of *MMP7* promoter in the south Asian (SAS) super population of 1000 genomes project. Pairwise LD values were plotted between common SNPs in the *MMP7* promoter using Haploview 4.2. D’: Coefficient of LD; LOD: log of the likelihood odds ratio.

### Occurrence of *MMP7* -181 SNP in Indian populations

The *MMP7* A-181G polymorphism was found to occur at a minor allele frequency (MAF) of 0.473 (∼72% of the study population), in the Chennai population comprising of 1517 hypertensive and normotensive individuals (Table 1). The study was replicated in 977 hypertensive and normotensive individuals from Chandigarh. The MAF of the *MMP7* A-181G SNP in the Chandigarh population was slightly lesser than that of the Chennai population, although still highly prevalent at 0.412 (64.7% of the study population). The genotypic frequencies in both populations were found to be in Hardy-Weinberg equilibrium (Chennai population: X^2^ p=0.99 and Chandigarh population: X^2^ p=0.37) (Table 1).

**Table 1.** Occurrence of *MMP7*-181A/G polymorphism in Indian populations

| ***MMP7***<br>**promoter**<br>**genotype** | **Occurrence in Chennai population** |  |  | **Occurrence in Chandigarh population** |  |  |
| --- | --- | --- | --- | --- | --- | --- |
|  | HTN<br>(n = 862) | Controls<br>(n = 655) | Total<br>(n = 1517) | HTN<br>(n = 519) | Controls<br>(n = 458) | Total<br>(n = 977) |
| A/A | 215 | 207 | 422 | 164 | 180 | 344 |
| A/G | 473 | 283 | 756 | 266 | 194 | 460 |
| G/G | 174 | 165 | 339 | 89 | 84 | 173 |
| Minor Allele<br>Frequency | 0.476 | 0.467 | 0.473 | 0.428 | 0.395 | 0.412 |
| H-W<br>p value | 0.991 |  |  | 0.372 |  |  |

Abbreviations: HTN, Hypertensive; H-W, Hardy-Weinberg equilibrium.

### *MMP7* -181AG genotype is associated with hypertension in Indian populations

Logistic regression analysis was carried out to study the relative risk contributed by the variant G allele towards hypertension after dividing the Chennai and Chandigarh populations into hypertensive cases and normotensive controls by both genotypic (AA *vs* AG and AA *vs* GG) and dominant (AA *vs* AG+GG) models. The -181AG heterozygous genotype was found to be significantly associated with hypertension risk when compared to -181AA genotype with an unadjusted odds ratio (OR) of 1.675 (95% Confidence Interval (CI) = 1.315-2.133; p=3.0×10^−5^) in Chennai population and 1.505 (95% CI = 1.136-1.993; p=0.004) in Chandigarh population (Table 2). The significance of these associations were retained even after adjusting for age, sex and body mass index [BMI] (Chennai population: OR = 1.641; 95% CI = 1.276-2.109; p=1.0×10^−4^and Chandigarh population: OR = 1.520; 95% CI = 1.106-2.090; p=0.01). Although the homozygous variant GG genotype did not show a statistically significant odds ratio, the dominant model (AG+GG) exhibited a significantly strong association with hypertension in both Chennai (adjusted OR = 1.408; 95% CI = 1.113-1.781; p=0.004) and Chandigarh (adjusted OR = 1.464; 95% CI = 1.085-1.977; p=0.013) populations (Table 2).

**Table 2.** Association of *MMP7*-181A/G polymorphism with hypertension risk in Indian populations*

| *MMP7*<br>promoter<br>genotype | Chennai population (n=1517) |  |  |  | Chandigarh population (n=977) |  |  |  |
| --- | --- | --- | --- | --- | --- | --- | --- | --- |
|  | Logistic Regression<br>(unadjusted) |  | Logistic Regression<br>(Age, Sex and BMI adjusted) |  | Logistic Regression<br>(unadjusted) |  | Logistic Regression<br>(Age and Sex adjusted) |  |
|  | OR (95% CI) | p value | OR (95% CI) | p value | OR (95% CI) | p value | OR (95% CI) | p value |
| A/A | 1(Ref) | - | 1(Ref) | - | 1(Ref) | - | 1(Ref) | - |
| A/G | **1.675 (1.315-2.133)** | **3 X 10<sup>-5</sup>** | **1.641 (1.276-2.109)** | **1 X 10<sup>-4</sup>** | **1.505 (1.136-1.993)** | **0.004** | **1.520 (1.106-2.090)** | **0.010** |
| G/G | 1.059 (0.796-1.411) | 0.693 | 1.017 (0.757-1.367) | 0.911 | 1.163 (0.807-1.676) | 0.418 | 1.314 (0.858-2.014) | 0.209 |
| A/G+G/G | **1.447 (1.154-1.815)** | **0.001** | **1.408 (1.113-1.781)** | **0.004** | **1.402 (1.077-1.824)** | **0.012** | **1.464 (1.085-1.977)** | **0.013** |

Abbreviations: OR, odds ratio; BMI, body mass index; CI, confidence interval

### Differential activities of *MMP7* -181A/G promoter-reporter constructs under basal conditions

To test the functional role of the -181A and -181G alleles in contributing to the promoter activity of the *MMP7* gene, *MMP7* promoter-reporter constructs harboring A and G alleles at the site of the polymorphism were generated in pGL3-basic vector. *MMP7*-181G and -181A promoter-reporter constructs transfected into cardiomyoblast (H9c2) and neuroblastoma cell lines (IMR-32, SH-SY5Y and N2a) showed differential promoter activities (Fig. 2). *MMP7*-181G construct consistently displayed higher promoter activity than the -181A construct in both cardiomyoblast H9c2 cell line (∼1.7 fold; p<0.01) and neuronal cell lines like IMR-32, SH-SY5Y and N2a (IMR-32: ∼1.4 fold; p<0.001; SH-SY5Y: ∼1.3 fold; p<0.05; N2a: ∼1.5 fold; p<0.05) suggesting that the variation is functional (Fig. 2).

**Figure 2.**
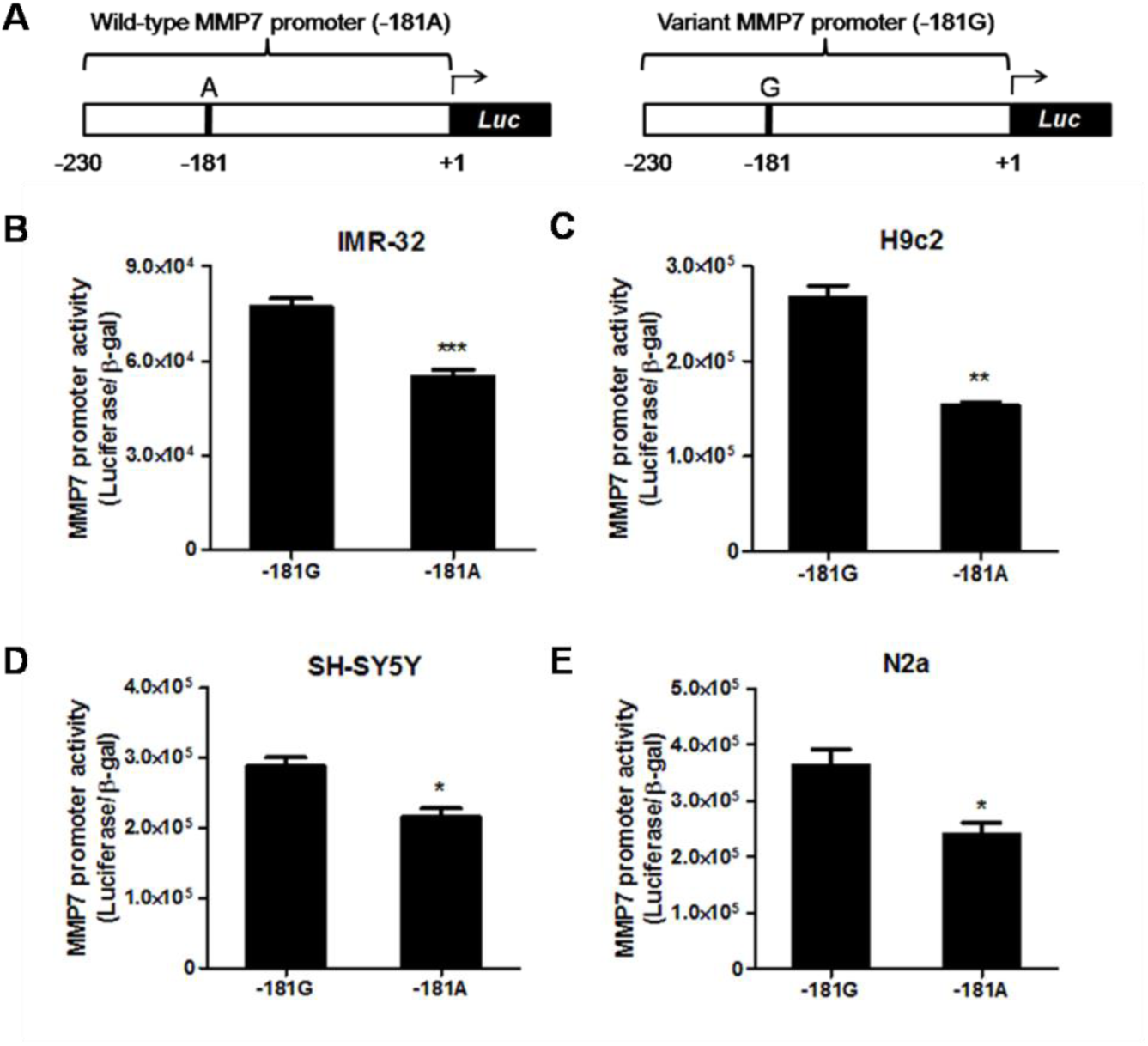
Differential promoter activities of *MMP7* promoter reporter constructs. **(A)** Schematic representation of the *MMP7* promoter reporter constructs cloned into pGL3-basic vector. **(B-E)** The *MMP7* promoter region harboring the -181A/G SNP from AA and GG individuals were cloned into pGL3-Basic plasmid. *MMP7* -181G and -181A promoter reporter constructs were transfected into rat cardiomyoblast H9c2 (B) neuroblastoma cell lines IMR-32 (C), SH-SY5Y (D) and N2a (E) along with β-galactosidase expression plasmid. Results are expressed as mean ± SEM of triplicate values of the ratio of luciferase/β-gal activity. *p<0.05, **p<0.01 and ***p<0.001 when compared to -181G construct.

### Computational analyses reveal differential interactions of CREB with *MMP7*-181A/G promoters

To probe for the possible differential interaction of transcription factors at -181bp position that could contribute to the differential activity of -181G and -181A alleles, computational prediction of transcription factors was carried out using ConSite, MatInspector and P-Match programs. The transcription factor CREB was predicted to bind to the -181G allele with higher affinity by both ConSite and P-Match programs. The Transfac position-weight matrix for CREB with binding scores for -181A and -181G alleles are shown in the Fig. S2. Following the computational prediction, further structure-based conformational and molecular dynamics simulation studies were carried out using models of CREB and the *MMP7* promoter.

The DP-Bind and PredictProtein tools predicted the region spanning amino acid residues 280-341 to have DNA binding affinity and hence the CREB1 homology model was generated for the region corresponding to residues 280-341 of CREB1. Models of *MMP7*-181A and *MMP7*-181G promoters were also generated and docking of the CREB1 structure to the promoter DNA structures followed by molecular dynamics simulations were carried out (Figs. 3A, S3 and Table S4).

**Figure 3.**
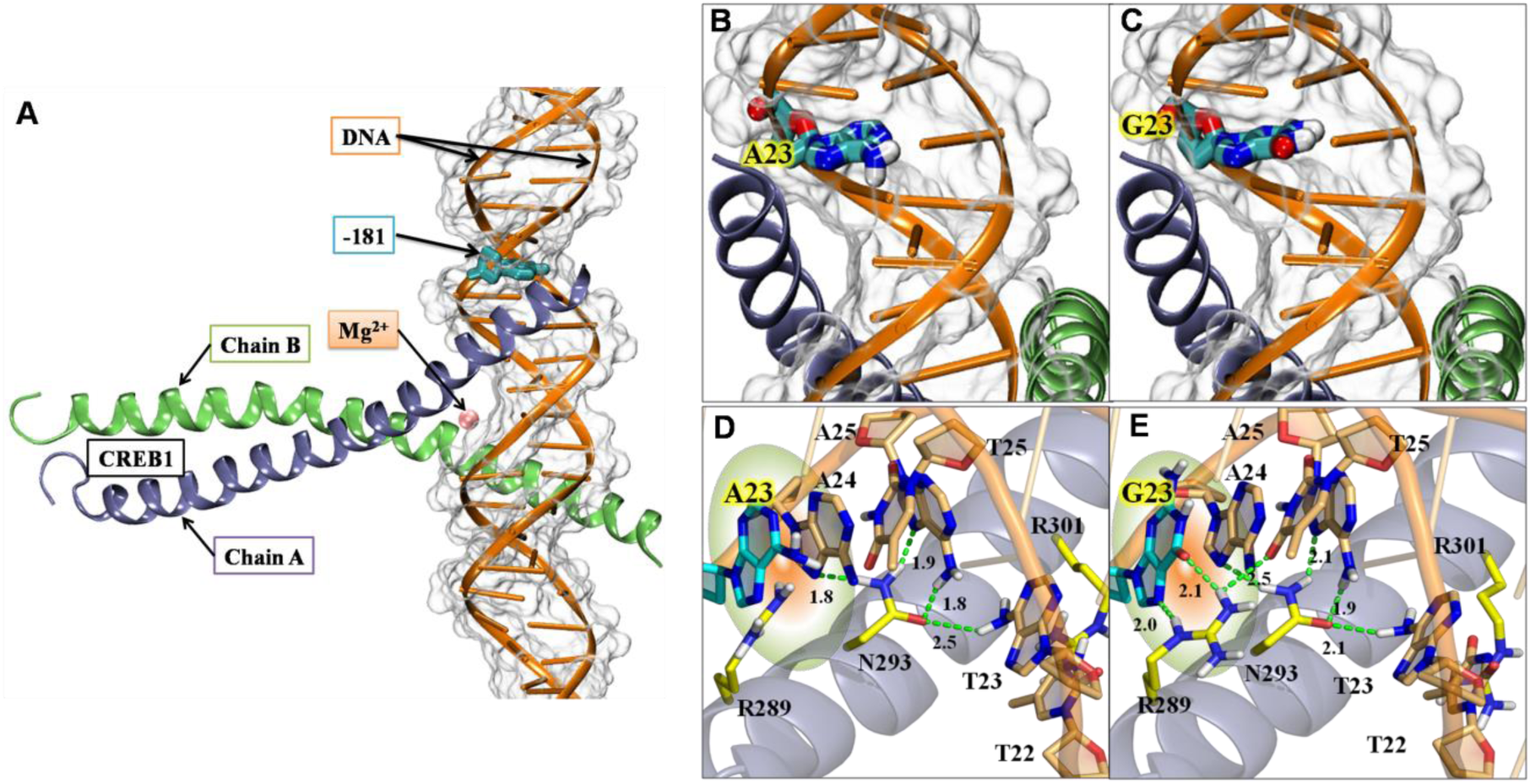
Schematic diagram of interactions of *MMP7*-promoter DNA with CREB1 transcription factor. **(A)** Representative energy minimized model of CREB1:wild-type *MMP7* complex rendered in new cartoon representation. DNA is shown in orange color; CREB1-chain A and CREB1-chain B are shown in ice-blue and green colors, respectively. DNA is shown in transparent white surface view as well. Position of the -181 bp nucleotide and Mg^2+^ has been indicated. **(B)** Positioning of the key nucleotide -181A (labelled as A23) in wild-type promoter. **(C)** Positioning of key nucleotide -181G (labeled as G23) in the mutant promoter. **(D)** and **(E)** Comparison of wild-type and mutant *MMP7*-promoter DNA interactions with CREB1 in enlarged view. Amino acids are indicated by single letter codes and based on their positions in the CREB1-chain A. Amino acids and nucleotides are rendered in licorice and colored atom wise; C: light orange, N: blue, O: red, H: white. Hydrogen bonds are shown in green dotted lines. The mutant promoter (E) involves several additional hydrogen bonds as compared to the wild-type promoter (D) suggesting stronger interactions with CREB1.

Since both complexes were stable till 40 ns in MD simulation, the averaged structure was extracted from the first 40 ns simulation to generate the comparable stable interaction map between *MMP7*-promoter DNA and CREB1. The interface sites were quantified in terms of residue-wise interaction energy, MM-GBSA (Molecular mechanics, the Generalized Born model and Solvent Accessibility) calculations and the lifetime occupancy of hydrogen bonds. The interaction pattern at the binding site of the *MMP7*-181A and *MMP7*-181G were quantified in terms of hydrogen bonds and hydrophobic contacts. In case of CREB1-chain A:*MMP7*-181A complex, N293 and R301 were the two main amino acids that participated in the interactions with the promoter DNA segment. The residue N293 formed total four hydrogen bonds (*viz.* one with A24 nucleotide at DNA-chain C, two with A25 nucleotide at DNA-chain C and one with T23 nucleotide at DNA-chain D) (Figs. 3B and 3D); the residue R301 formed one hydrogen bond with T22 nucleotide at DNA-chain D. On the other hand, the CREB1-chain A:*MMP7*-181G complex apart from having interactions similar to that of *MMP7*-181A complex, also established three additional hydrogen bonds with residue R289 (two hydrogen bonds with G23 nucleotide at DNA-chain C [i.e., the -181G variant nucleotide] and one hydrogen bond with T25 nucleotide at DNA-chain D) (Figs. 3C and 3E).

The 2D-interaction map of the *MMP7*-181A and *MMP7*-181G systems reflects the same interaction pattern (Fig. S4). The interaction map displays two regions of interactions for CREB1-chain A, between 22-26 nucleotide on *MMP7* DNA:chain C (i.e., -180 to -184 bp of *MMP7* promoter) and 21-22 nucleotide on *MMP7* DNA:chain D (i.e., -185 to -186 bp of *MMP7* promoter). Similarly, CREB1-chain B interacts with 29-31 nucleotide on *MMP7* DNA:chain C (i.e., -187 to -189 bp of *MMP7* promoter) and 15-18 nucleotide on *MMP7* DNA:chain D (i.e., -189 to -192 bp of *MMP7* promoter). Despite differences in the van der Waals contacts made by the amino acid residues of CREB1-chain A and CREB1-chain B with the *MMP7* wild-type and mutated promoters, the direct hydrogen bonds between the *MMP7*-181G nucleotide and R289 residue (identified as a hot-spot residue in case of CREB1:*MMP7*-181G complex alone but not in CREB1:*MMP7*-181A complex) could facilitate the favorable binding of CREB to the *MMP7*-181G promoter with higher affinity over *MMP7*-181A promoter.

### Experimental evidence for interactions of CREB with *MMP7*-181A/G promoters

In view of the prediction of enhanced interactions of CREB with the *MMP7*-181G allele (Figs. 3 and S4), experimental validation of differential activation of *MMP7*-181A/G promoters by CREB was carried out. Over-expression of CREB increased the promoter activity of the *MMP7* promoter-reporter constructs in a dose-dependent manner with the-181G construct exhibiting significantly higher promoter activity than the -181A construct in IMR-32 as well as H9c2 cells (IMR-32: one-way ANOVA F=162.9, p<0.0001; H9c2: one-way ANOVA F=43.3, p<0.0001) (Figs. 4A and 4B). Over-expression of CREB was confirmed by Western blotting. In corroboration, co-transfection of dominant negative KCREB plasmid lead to a highly significant dose-dependent decrease (at least 60%) in the promoter activity of *MMP7*-181G construct in both IMR-32 and H9c2 cells (IMR-32: one-way ANOVA F=13.72, p<0.0001; H9c2: one-way ANOVA F=11.45, p<0.001) (Figs. 4C and 4D). The -181A construct did not show any change in promoter activity in response to KCREB co-transfection. Down-regulation studies using CREB siRNA oligo also resulted in ∼45% and ∼60% reduction in promoter activity of -181G construct when compared to the negative control oligo in IMR-32 and H9c2 cells respectively (IMR-32: one-way ANOVA F=7.13, p<0.05; H9c2: one-way ANOVA F=79.02, p<0.0001) when compared to the -181A construct (Figs. 5A and 5B). Down-regulation of CREB was confirmed by Western blotting.

**Figure 4.**
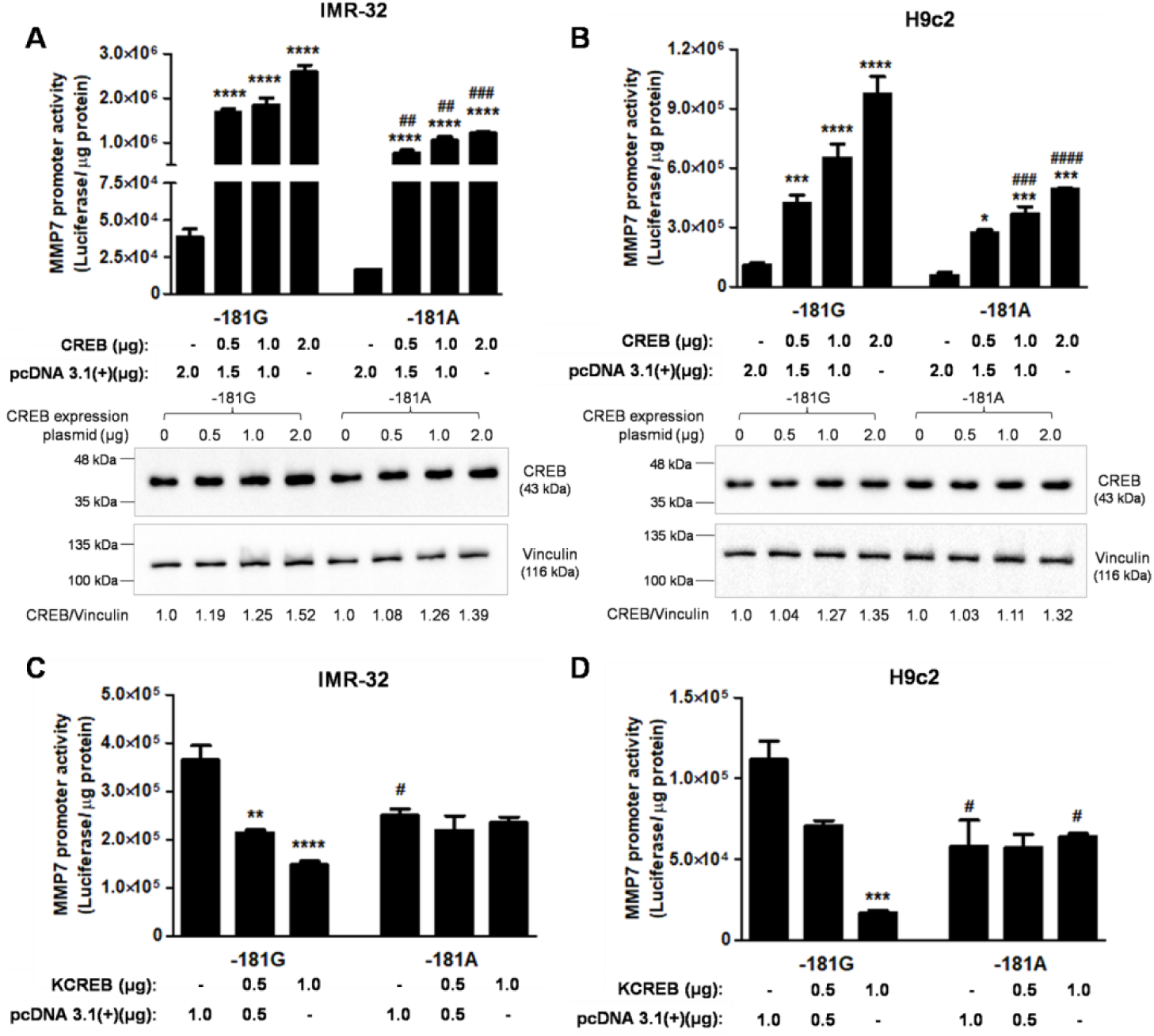
Role of CREB in augmentation of *MMP7* promoter activity. IMR-32 (A) and H9c2 (B) cells were transiently transfected with *MMP7* -181G and -181A promoter-reporter constructs and increasing doses of CREB expression plasmid. pcDNA 3.1(+) was used as balancing plasmid. Assays were performed 30 hours after transfection and results expressed as mean ± SEM of triplicate values of the ratio of luciferase activity/μg protein. *p<0.05, **p<0.01 and ***p<0.001 when compared to basal activity of the corresponding construct. ##p<0.01, ###p<0.001 and ####p<0.0001 as compared to corresponding dose in case of the *MMP7*-181G construct. Over-expression of CREB was confirmed by western blotting. **(C)** and **(D)** Dominant negative KCREB diminishes the promoter activity of *MMP7*-181G promoter. IMR-32 (C) and H9c2 (D) cells were transiently transfected with *MMP7* promoter-reporter constructs and increasing quantities of KCREB plasmid. pcDNA3.1(+) was used as balancing plasmid. Assays were performed 30 hours after transfection and results expressed as mean ± SEM of triplicate values of the ratio of luciferase activity/μg protein. **p<0.01, ***p<0.001 and ****p<0.0001 when compared to basal activity of the corresponding construct. #p<0.05 as compared to corresponding dose in case of the *MMP7*-181G construct.

**Figure 5.**
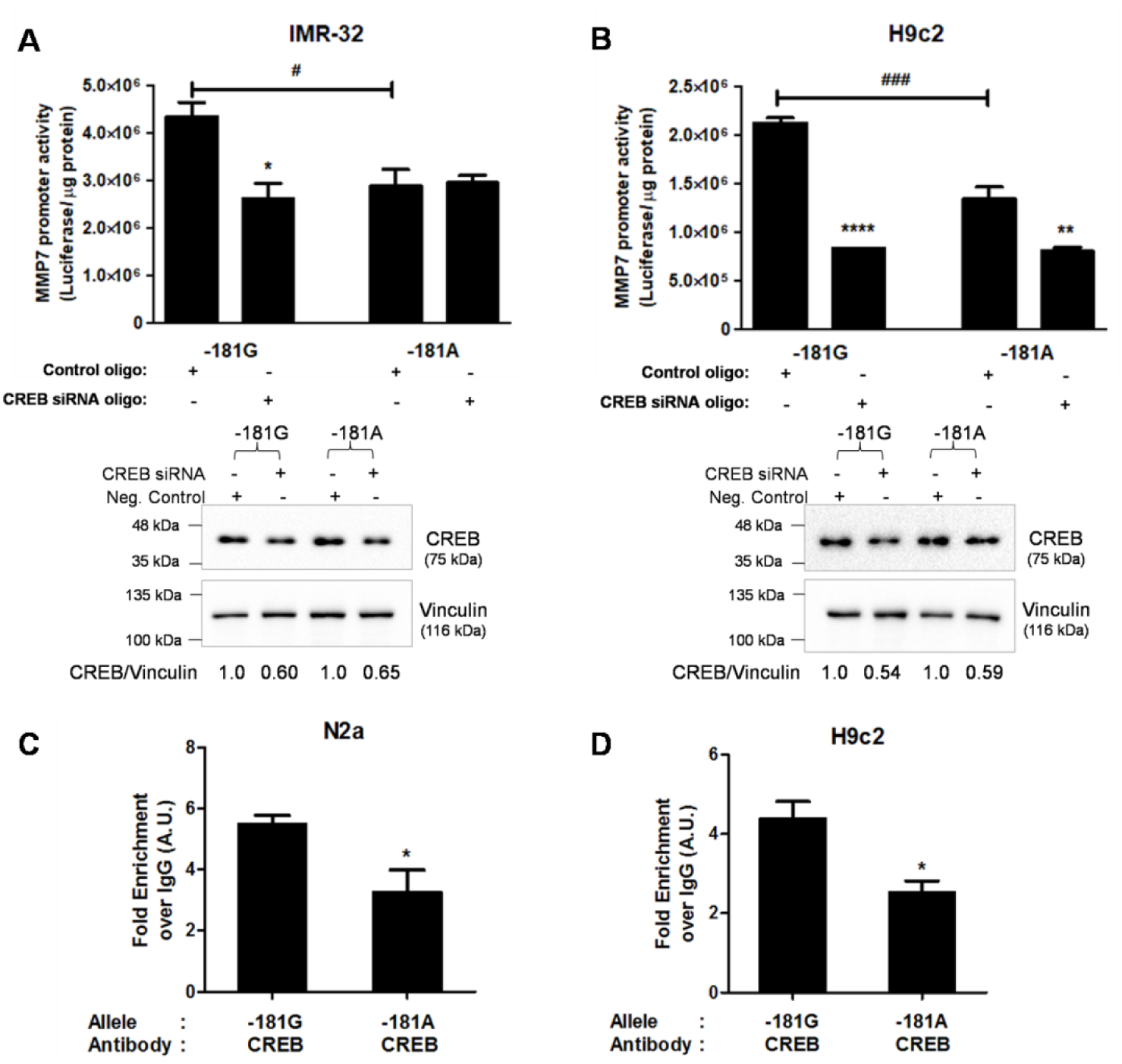
Enhanced interactions of CREB with *MMP7* -181G allele *in vivo* in the context of chromatin. **(A)** and **(B)** siRNA mediated knock-down of CREB decreases the *MMP7*-181G promoter activity. IMR-32 (A) and H9c2 (B) cells were transfected with *MMP7* -181G or -181A constructs along with control siRNA oligo or CREB siRNA oligo. Assays were performed after 48 hours of transfection and results expressed as mean ± SEM of triplicate values of the ratio of luciferase activity/μg protein. *p<0.05, **p<0.01 and ****p<0.0001 when compared to the basal activity of the corresponding construct. #p<0.05 and ### p<0.001 as compared to basal activity of *MMP7*-181G construct. Down-regulation of CREB was confirmed by western blotting. **(C)** and **(D)** Interaction of CREB with -181G/A alleles of *MMP7* promoter. Binding of CREB to -181G and -181A alleles of *MMP7* promoter. Chromatin immunoprecipitation of N2a (C) and H9c2 (D) cells transfected with -181G or -181A construct was carried out using antibody against CREB/control IgG. Immunoprecipitated chromatin was subjected to qPCR. Fold enrichment in case of CREB antibody over IgG control is shown. CREB displayed higher binding affinity towards -181G promoter when compared to the -181A promoter.

Further, chromatin immunoprecipitation (ChIP) assays were carried out to study the interaction and preferential binding of CREB with the *MMP7*-181G promoter in the context of chromatin. Quantitative PCR for purified, immunoprecipitated chromatin from N2a and H9c2 cells transfected with *MMP7*-181G and -181A promoter constructs revealed that enrichment with -181G allele of the *MMP7* promoter was ∼1.7-and ∼1.8-fold higher following immunoprecipitation with CREB when compared with -181A allele (p<0.05) (Figs. 5C and 5D). These results strongly suggest that the higher promoter activity of *MMP7*-181G construct could be attributed to higher binding affinity of CREB towards the -181G allele.

### Enhanced response of *MMP7*-181G promoter to epinephrine: Crucial role for CREB

Since elevated levels of catecholamines are associated with hypertension we probed the effect of epinephrine and the concomitant role of CREB, if any, on *MMP7*-181G and -181A promoter activities. Indeed, epinephrine treatment augmented the promoter activity of *MMP7*-181G construct in a dose-dependent manner up to ∼2.5-fold and ∼2.2-fold, respectively, in IMR-32 and H9c2 cells (IMR-32: one-way ANOVA F=22.03, p<0.0001; H9c2: one-way ANOVA F=17.10, p<0.0001) (Figs. 6A and 6B); on the other hand, *MMP7*-181A construct did not show such significant increase in promoter activity. Consistently, epinephrine treatment enhanced ∼4.4-and ∼12.4-fold (when normalized to vinculin) increase in phospho-CREB levels in IMR-32 and H9c2 cells, respectively (Figs. 6C and 6D). Further, to test if CREB mediated the activation of *MMP7*-181G promoter in response to epinephrine, IMR-32 and H9c2 cells were co-transfected with KCREB and *MMP7*-181G or -181A promoter-reporter constructs followed by epinephrine treatment. Epinephrine, which augmented the promoter activity of *MMP7*-181G construct significantly failed to evoke a similar response in the KCREB co-transfected condition (IMR-32: two-way ANOVA genotype effect: F=56.05, p<0.0001; treatment effect: F=15.30, p<0.0001; H9c2: two-way ANOVA genotype effect: F=16.50, p<0.001; treatment effect: F=17.58, p<0.0001). The -181A construct did not show any significant difference in promoter activity upon epinephrine treatment either in the presence or absence of KCREB (Figs. 6E and 6F).

**Figure 6.**
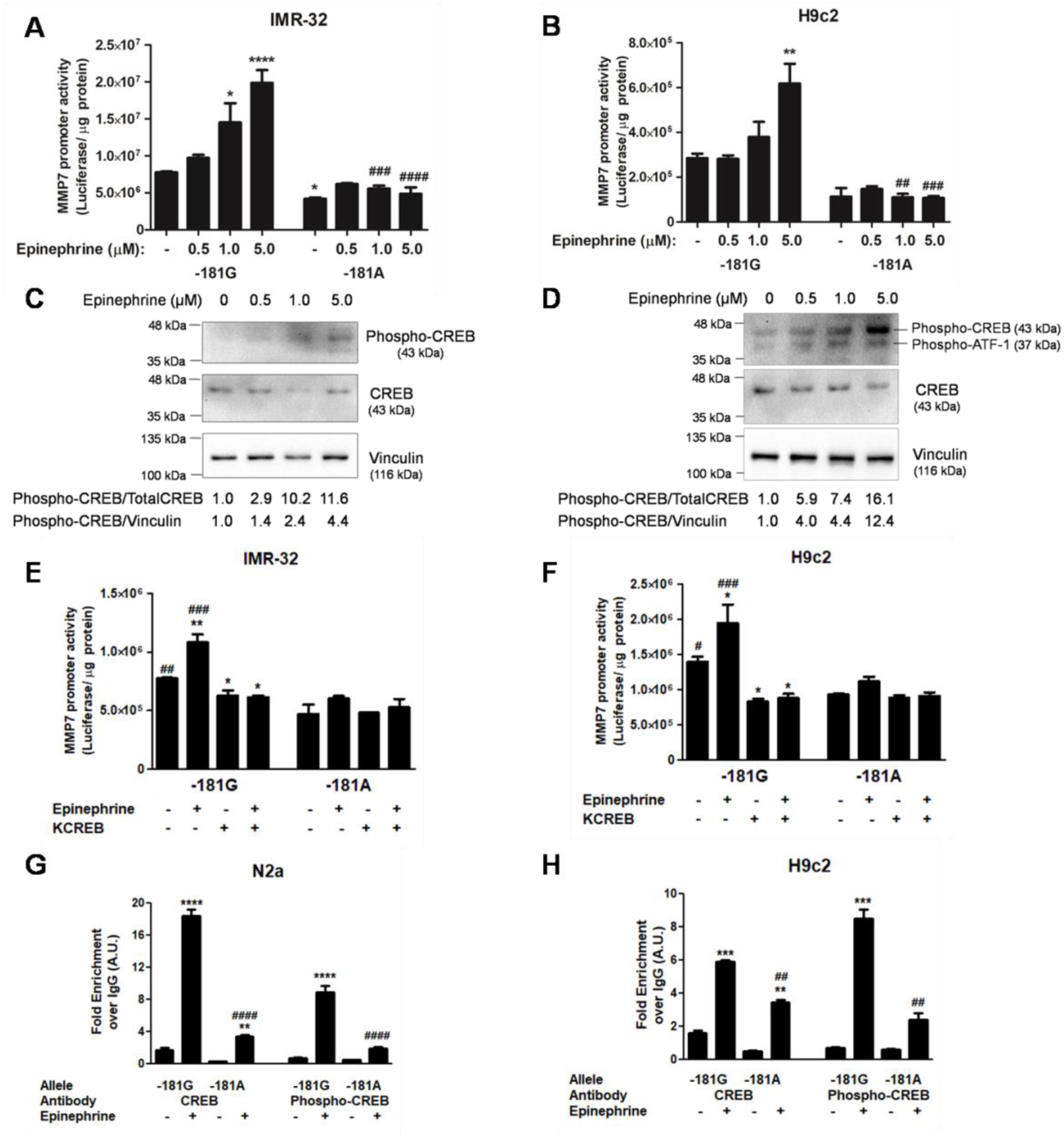
Epinephrine induced activation of *MMP7* -181G promoter via CREB. IMR-32 (A) and H9c2 (B) cells were transfected with *MMP7* promoter -181G and -181A constructs and treated with increasing doses of epinephrine for 6 hrs. Results are mean ± SEM of triplicate values of luciferase activity/μg protein. *p<0.05, **p<0.01 and ****p<0.0001 when compared to respective basal condition and ##p<0.01, ###p<0.001 and ####p<0.0001 when compared to the corresponding treatment in case of -181G construct. **(C)** and **(D)** Effect of epinephrine treatment on CREB expression. IMR-32 (C) and H9c2 (D) cells were treated with increasing doses of epinephrine for 6 hrs and western blot analysis of total proteins probing for phospho-CREB, CREB and Vinculin was carried out. **(E)** and **(F)** Role of CREB in allele specific activation of *MMP7* promoter -181G by epinephrine. *MMP7* promoter -181G and -181A constructs were transfected into IMR-32 (E) and H9c2 (F) cells with or without dominant negative KCREB plasmid followed by treatment with epinephrine (5 μM) for 6 hrs. Results are expressed as mean ± SEM of triplicate values of luciferase activity/μg protein. *p<0.05 and **p<0.01 when compared to basal condition and #p<0.05, ##p<0.01 and ###p<0.001 when compared to the corresponding condition in -181A construct. **(G)** and **(H)** ChIP assay of N2a and H9c2 cells transfected with *MMP7* promoter -181G or -181A construct, with or without epinephrine treatment was carried out using antibody against CREB, phospho-CREB and IgG. Immunoprecipitated chromatin was subjected to qPCR. Fold enrichment in signal obtained in case of CREB and phospho-CREB antibodies over IgG control is shown. **p<0.01, ***p<0.001 and ****p<0.0001 when compared to fold enrichment in respective untreated condition and ##p<0.01 and ####p<0.0001 when compared to the corresponding treatment in case of -181G construct.

In ChIP assays using N2a and H9c2 cells treated with epinephrine following transfection of the *MMP7* promoter constructs, the -181G allele displayed significantly higher fold enrichment with CREB (N2a: ∼5.0-fold; H9c2: ∼1.8-fold) and phospho-CREB (N2a: ∼4.5-fold; H9c2: ∼3.5-fold) as compared to the -181A allele (N2a: one-way ANOVA F=223.9, p<0.0001; H9c2: one-way ANOVA F=128.1, p<0.0001) (Figs. 6G and 6H). Thus, the epinephrine induced activation of *MMP7*-181G promoter activity appears to be strongly mediated by CREB.

### Hypoxic stress activates *MMP7*-181G promoter via CREB

Since hypoxia is known to increase blood pressure and is a major contributor to cardiac pathophysiology, the effect of hypoxia on the promoter activity of *MMP7*-181G and -181A constructs and the role of CREB towards the same was studied. IMR-32 and H9c2 cells were subjected to hypoxia after transfection with *MMP7*-181G and -181A promoter constructs for 6 hrs with/without CREB and KCREB co-transfection. The basal promoter activity of *MMP7*-181G promoter was enhanced significantly in response to hypoxia in both IMR-32 and H9c2 cells (IMR-32: ∼1.7-fold; H9c2: ∼2.3-fold); however, no significant increase was observed in case of the -181A construct (Figs. 7A and 7B). In the CREB co-transfected condition hypoxia treatment further increased the promoter activity of the *MMP7*-181G construct (∼5.7-fold in IMR-32 and ∼7.5-fold in H9c2) when compared to *MMP7*-181A construct (IMR-32: two-way ANOVA genotype effect: F=39.7, p<0.0001; treatment effect: F=267.8, p<0.0001; H9c2: two-way ANOVA genotype effect: F=30.97, p<0.0001; treatment effect: F=73.01, p<0.0001). Similarly, hypoxia resulted in only a modest increase in promoter activity in the KCREB co-transfected cells. Along with HIF-1α over-expression (IMR-32: ∼6.6-fold; H9c2: ∼7.9-fold), CREB (IMR-32: ∼1.4-fold; H9c2: ∼1.8-fold), and phospho-CREB (IMR-32: ∼1.5-fold; H9c2: ∼1.3-fold) levels were also found to be elevated under hypoxia by Western blot analysis (Figs. 7C and 7D).

**Figure 7.**
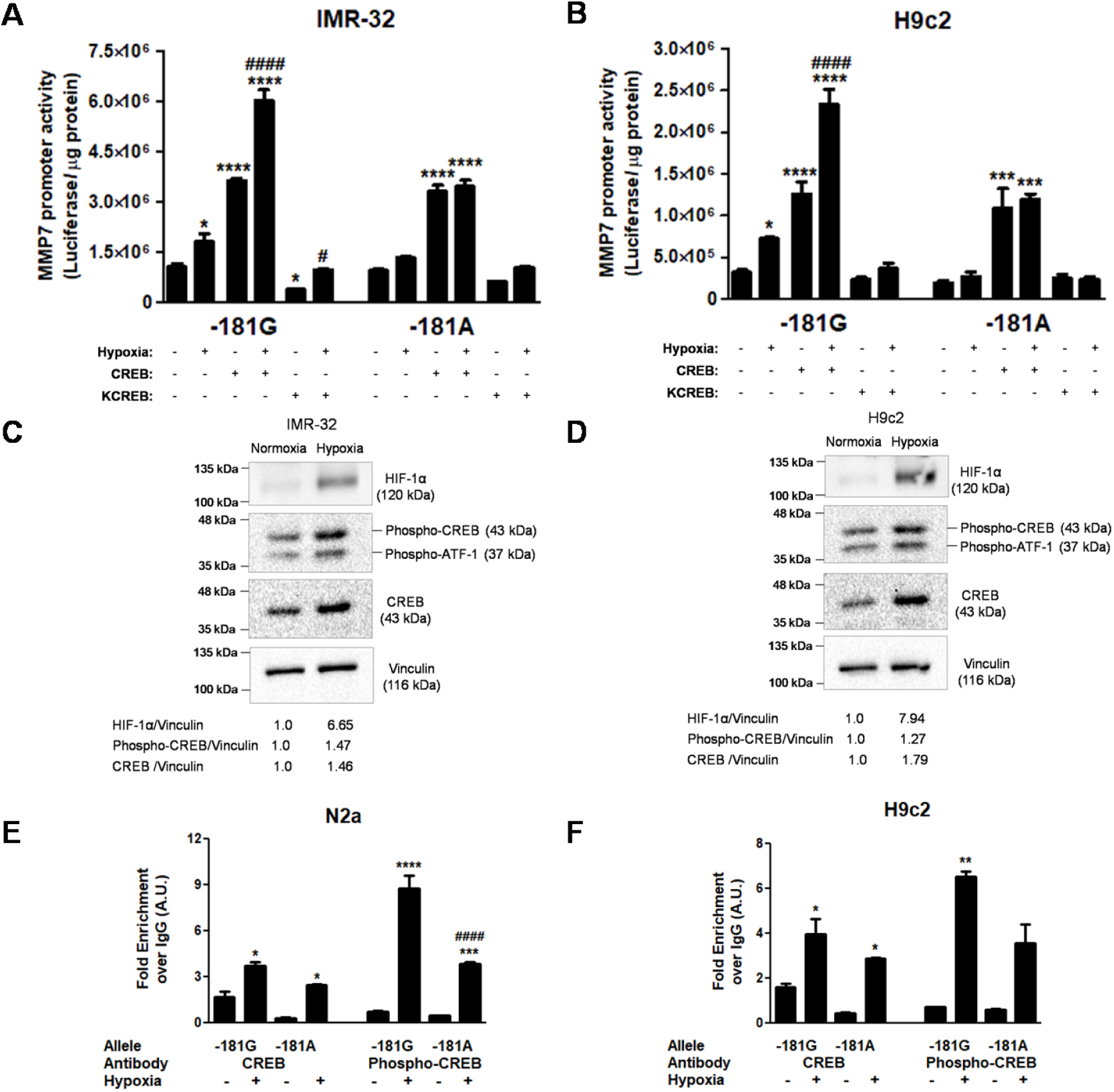
Effect of hypoxia on *MMP7* promoter: crucial role for CREB. **(A)** and **(B)** Role of CREB in allele specific activation of *MMP7* -181G promoter in response to hypoxia. *MMP7* -181G and -181A promoter reporter constructs were transfected into IMR-32 (A) and H9c2 (B) cells with/without CREB and KCREB expression plasmid and subjected to 12 hrs of hypoxia. Results are expressed as mean ± SEM of triplicate values of luciferase activity/μg protein, *p<0.05, ***p<0.001 and ****p<0.0001 when compared to basal activity of the respective construct and #p<0.05 and ####p<0.0001 when compared to the corresponding normoxic condition. **(C)** and **(D)** Effect of hypoxia on CREB expression. IMR-32 (C) and H9c2 (D) cells were subjected to hypoxia for 12 hrs and western blotting of the total proteins was carried out probing for HIF1-α, CREB, phospho-CREB and vinculin. **(E)** and **(F)** Effect of hypoxia on the binding of CREB to *MMP7* -181G and -181A alleles. Chromatin immunoprecipitation of N2a and H9c2 cells transfected with *MMP7* -181G or -181A promoter construct, with/without exposure to hypoxia was carried out using antibody against CREB, phospho-CREB and pre-immune IgG. Immunoprecipitated chromatin was subjected to qPCR and fold enrichment in signal obtained in case of CREB/phospho-CREB antibody over pre-immune IgG is shown. *p<0.05 **p<0.01, ***p<0.001 and ****p<0.0001 when compared to fold enrichment in respective untreated condition and ####p<0.0001 as compared to the corresponding treatment in case of *MMP7* -181G promoter construct.

Further, the interaction of CREB and phospho-CREB with the -181G and -181A alleles of *MMP7* promoter under hypoxia was probed in N2a and H9c2 cells transfected with *MMP7* promoter-reporter constructs. The fold enrichment for CREB (N2a: ∼1.5-fold; H9c2: ∼1.4-fold) and phospho-CREB (N2a: ∼2.3-fold; H9c2: ∼1.9-fold) observed in case of -181G allele in response to hypoxia was significantly higher (Figs. 7E and 7F) than the -181A allele (N2a: one-way ANOVA F=63.7, p<0.0001; H9c2: one-way ANOVA F=28.0, p<0.0001).

### *MMP7*-181G promoter results in higher MMP7 levels *in vitro* and *in vivo*

By generating recombinant plasmids wherein hMMP7 cDNA was placed under the control of *MMP7*-181G and *MMP7*-181A promoters we probed whether the differential activities *MMP7*-181G and -181A result in altered MMP7 levels *in vitro* (Fig. 8A). These constructs were transfected into N2a (Figs. 8B and 8C) and H9c2 (Figs. 8D and 8E) cells and Western blot analysis of cell lysates revealed significantly higher levels of MMP7 in case of -181G promoter-driven *MMP7* cDNA (N2a: ∼1.6-fold; p<0.0001; H9c2: ∼1.4-fold; p<0.05) as compared to -181A promoter driven MMP7 cDNA suggesting that in the genomic context the *MMP7*-181G promoter is likely to be more active than the -181A promoter.

**Figure 8.**
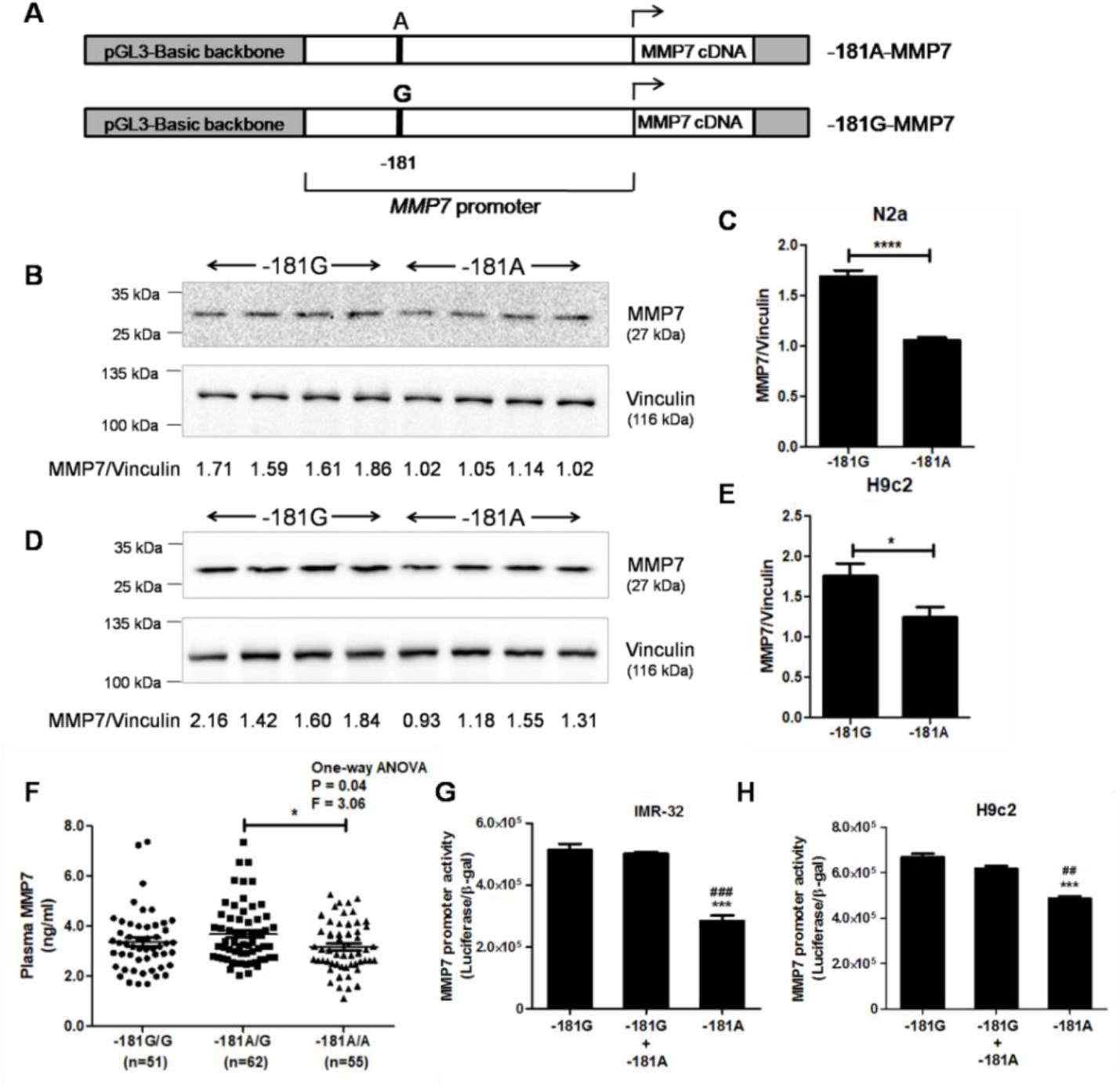
*MMP7* -181G promoter confers higher expression of MMP7 *in vitro*. **(A)** Schematic representation of the *MMP7* promoter-cDNA constructs. The *MMP7* cDNA was subcloned in frame into the -181G and -181A promoter reporter constructs replacing the luciferase cDNA. **(B)** and **(C)** *In vitro* expression of MMP7 under the influence of *MMP7* -181G and -181 A promoters in N2a cells. N2a cells were transfected with -181G-MMP7 and -181A-MMP7 constructs and western blotting (B) of total proteins was carried out to probe for MMP7 and vinculin. Fold difference (MMP7/vinculin) is indicated (C). **(D)** and **(E)** *In vitro* expression of MMP7 under the influence of *MMP7* -181G and -181A promoters in H9c2 cells. H9c2 cells were transfected with -181G-MMP7 and -181A-MMP7 constructs and western blotting (D) of total proteins was carried out to probe for MMP7 and vinculin. Fold difference (MMP7/vinculin) is indicated (E). **(F)** Plasma MMP7 levels in -181A/A, -181A/G and -181G/G individuals as measured by ELISA. *MMP7* -181A/G individuals presented with significantly higher MMP7 levels. **(G)** and **(H)** *MMP7*-181A/G promoter activities in diploid combinations *in cella*. IMR-32 (G) and H9c2 (H) cells were transfected with 1 μg each of *MMP7* -181G or -181A or 500 ng each of -181G and -181A constructs. Results are mean ± SEM of triplicate values of luciferase/β-gal activity. ***p<0.001 as compared to -181G construct and ##p<0.01, ###p<0.001 as compared to -181G+-181A construct.

Further, plasma MMP7 levels in a section of the study population after stratification into *MMP7* promoter genotypes revealed that *MMP7*-181A/G individuals displayed higher plasma MMP7 levels (Fig. 8F) than -181A/A and -181G/G individuals (one-way ANOVA F=3.06, p<0.05). Although plasma MMP7 levels were modestly higher in individuals of -181G/G genotype as compared to those with -181A/A genotype, the difference was not statistically significant. Thus, the plasma levels of MMP7 follow the same trend of association displayed by the -181A/G individuals in the logistic regression analysis towards hypertension risk (Table 2).

### Activities of *MMP7* promoter constructs in diploid combinations *in cella*

In view of the association of *MMP7*-181A/G genotype with hypertension risk as well as higher MMP7 levels *in vivo* we sought to test whether this observation could also be demonstrated in transfected cultured cells. We transfected IMR-32 and H9c2 cells with -181G or -181A promoter-reporter plasmids in three diploid combinations: -181G/G (i.e., only -181G construct to mimic the homozygous variant), -181G/A (i.e., equimolar amounts of -181G and -181A constructs to mimic the heterozygous condition) and -181A/A (i.e., only -181A construct to mimic the wild-type condition). Interestingly, both the -181G and the -181G/A transfected conditions resulted in almost similar extent of elevated promoter activity (∼1.8-fold and ∼1.75-fold, respectively, in IMR-32 cells; ∼1.4-fold and ∼1.3-fold, respectively, in H9c2 cells) than the -181A construct alone (IMR-32: one-way ANOVA F=63.74, p<0.0001; and H9c2: one-way ANOVA F=51.37, p<0.001) (Figs. 8G and 8H). These results suggest that the *MMP7*-181G allele may act in a dominant manner while impacting the promoter activity.

### Association of *MMP7* -181AG genotype with blood pressure

To understand the potential associations of these genotypes with traits influencing cardiovascular physiology inferential statistics was carried out using the phenotypic data for demographic/physiological/biochemical parameters of the study population after stratifying them on the basis of their *MMP7*-181 promoter genotype. We observed higher mean arterial pressure [MAP] (∼2-3 mm Hg) and diastolic blood pressure [DBP] (∼2-3 mm Hg) in individuals of *MMP7* -181AG genotype than -181AA or -181GG individuals in both Chennai (MAP: one-way ANOVA F=3.17, p<0.05; DBP: one-way ANOVA F=3.70, p<0.05) and Chandigarh (MAP: one-way ANOVA F=3.06, p<0.05; DBP: one-way ANOVA F=3.29, p<0.05) populations (Figs. 9A and 9B). The MAP and DBP levels in these subjects positively correlated with plasma MMP7 levels (MAP: Pearson r=0.21, p<0.01; DBP: Pearson r=0.29, p<0.001) (Figs. 9C and 9D). *MMP7*-181A/G individuals also displayed a trend towards higher systolic blood pressure (∼3 mm Hg), although the difference was not statistically significant (Fig. S5). However, a modest positive correlation was observed with plasma MMP7 levels in these individuals (Pearson r=0.17, p<0.05) (Fig. S5).

**Figure 9.**
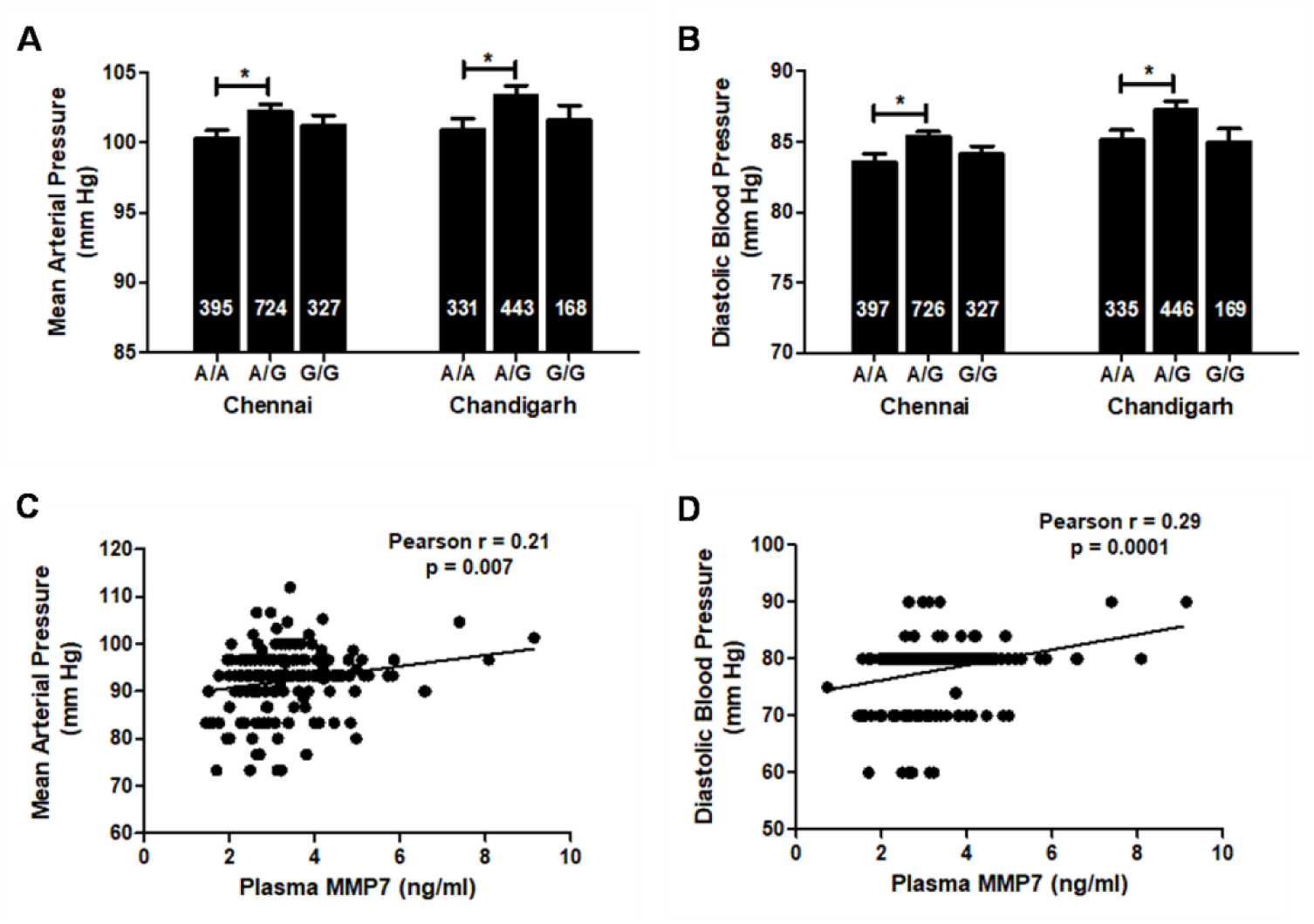
Allele-specific association of *MMP7*-181 promoter genotypes with blood pressure in Indian populations. **(A)** and **(B)** *MMP7*-181A/G individuals display higher mean arterial pressure (A) and diastolic blood pressure (B) in Chennai and Chandigarh populations. **(C)** and **(D)** Correlation of diastolic blood pressure (C) and mean arterial pressure (D) with MMP7 levels. MMP7 levels showed significant positive correlation with systolic, diastolic blood pressure and mean arterial pressure. Pearson r and p values for the correlations are indicated.

## DISCUSSION

### MMPs and their role in cardiovascular diseases

Hypertension, a major risk factor for cardiovascular disease is characterized by several complex pathophysiological mechanisms including cardiac remodelling.^22^ The remodelling which occurs initially as an adaptive response results in increased/altered ECM content ultimately leading to cardiovascular dysfunction.^23^ MMPs play a crucial role in cardiac remodelling by degrading extracellular matrix components which provide structural and mechanical support to the vasculature.^24^ An imbalance between the activities and levels of MMPs and their inhibitors (TIMPs) has been implicated in the pathogenesis of hypertension and cardiovascular diseases.^25^ Elevated MMP levels have been shown to increase blood pressure via cleavage of pharmacological receptors (e.g., β2-adrenergic receptor, epidermal growth factor receptor) and vasoactive peptides (e.g., endothelin, adrenomedullin).^8^

MMP7, the smallest known member of the MMP family is known to cleave a wide range of substrates including ECM components, vasoactive ligands, growth factor receptors, pro-inflammatory molecules and other MMPs.^26^ Knock-down of *MMP7* in spontaneous hypertension rat model attenuated hypertension and stopped the development of cardiac hypertrophy.^9^ MMP7 was also shown to be involved in early stages of agonist induced hypertension where it served as a transcriptional regulator of MMP2 in Ang-II induced hypertension. The study revealed that inhibition of MMP2 prevented only blood pressure elevation whereas knock-down of MMP7 and TACE (TNF-α converting enzyme) prevented hypertension as well as the development of cardiac hypertrophy.^10^ Since expression of MMPs including MMP7 is known to be tightly regulated at transcriptional level, polymorphisms in the promoter region may affect the gene expression by altering the binding affinities of transcription factors binding to the transcription factor binding sites in the promoter. Such functional polymorphisms could potentially increase/decrease susceptibility or genetic predisposition to pathophysiological phenotypes.^11^ Two polymorphisms in the MMP7 promoter (rs11568818:A-181G and rs11568819:C-153T) have been shown to be associated with the risk of different types of cancers, coronary artery disease^7, 11^ but no studies have reported a direct association of these variants with hypertension risk. Hence, we sought out to study the association of *MMP7* promoter variation with hypertension risk if any in two geographically distinct Indian populations using a systematic case-control approach followed by functional characterization of the associated variation.

### Occurrence and association of *MMP7*A-181G polymorphism with hypertension in geographically distinct Indian populations

We first performed pair-wise linkage disequilibrium analysis of common SNPs in the *MMP7* upstream promoter region (Fig. 1) and identified rs11568818 (*MMP7* A-181G) as a tag SNP, which was used for further genotyping and functional characterization. The *MMP7* A-181G polymorphism was found to occur at a frequency of 0.473 in the primary Chennai population and at a frequency of about 0.412 in the replication population from Chandigarh (Table 1). The polymorphism occurred at a frequency similar to that observed in the European, south Asian and African super populations (EUR: 0.44, SAS: 0.43, AFR: 0.45) of the 1000 Genomes project. Interestingly, the polymorphism occurred at a lesser frequency of 0.34 in the American super population and was least frequent in the east Asian super population with a mere frequency of 0.085 suggesting an ethnicity dependent occurrence of the SNP in different populations.

Next, we probed for association of the *MMP7* A-181G polymorphism with hypertension risk. The -181 A/G genotype conferred at least ∼1.5-fold higher risk of hypertension in both the Chennai population and in the replication population from Chandigarh with the dominant model also displaying a highly significant association with hypertension risk (Table 2). A multitude of studies have demonstrated the association *MMP7* A-181G polymorphism with several types of cancers, however only limited reports are available characterizing the association of this SNP in cardiovascular disorders. In a meta-analysis of 25 studies comprising of 6392 cases and 7665 controls that suggested a higher overall cancer risk for the *MMP7* A-181G polymorphism especially among Asian populations, the GG genotype was particularly associated with increased risk of colorectal cancer.^27^ The heterozygous A/G genotype also enhanced susceptibility to grade II astrocytoma by 2-fold while G/G genotype increased susceptibility of grade II to IV astrocytoma by 3-folds.^28^ The *MMP7* -181G allele was also found to significantly increase the risk of esophageal carcinoma, gastric cardiac adenocarcinoma, non-small cell lung cancer, ovarian cancer.^7, 29^ In a study involving a north Indian population, an increased risk of cervical cancer was conferred on to individuals with -181GG genotype (OR=1.94) or -181G allele (OR=1.37).^30^ Recently, we showed association of *MMP7* -181GG genotype with increased gastric cancer susceptibility (OR=1.9; p=0.02).^5^ Although no independent association of *MMP7* A-181G polymorphism was observed with left ventricular dysfunction in CAD patients in a study conducted in north Indian population^31^, high order gene-gene interaction analysis using classification and regression tree (CART) and multifactor dimensionality reduction (MDR) with four-factor interaction model predicted that individuals carrying the combination of AT1 1166AC+CC, NFKB1 -94 ATTG Ins/Ins, *MMP7* -181AG+GG, and MMP9 668RQ+QQ genotypes displayed a significantly higher risk for LVD (adjusted OR=8.14; p=0.003).^32^ To follow up this first report on the association of *MMP7* A-181G SNP with hypertension risk it would be interesting to carry out systematic large-scale studies on the potential association of this genetic variation with related cardiovascular disease states in different human populations.

### Allele-specific effect of CREB in activation of *MMP7* -181G promoter in basal and pathophysiological conditions

Transient transfections of *MMP7*-181A and -181G promoters revealed that the -181G promoter consistently displayed higher promoter activity than the -181A promoter across different cell lines suggesting that the variation was functional (Fig. 2). Next, we probed the molecular mechanism behind the observed higher promoter activity of the -181G promoter. Detailed computational analyses including structure-based conformational and molecular dynamics simulation studies affirmed stronger interaction of CREB transcription factor with the -181G promoter (Figs. 3 and S4). Co-transfection experiments in H9c2 and IMR-32 cells with CREB over-expression, dominant negative constructs and siRNA mediated down-regulation studies corroborated the results from the computational analysis; the -181G promoter construct displayed a higher promoter activity in response to CREB over-expression (Figs. 4A and 4B) while exhibiting a greater reduction in promoter activity in response to CREB knock-down experiments (Figs. 4 and 5) as compared to the -181A promoter construct. Chromatin immunoprecipitation studies further confirmed enhanced interaction of CREB with -181G promoter as compared to -181A promoter (Fig. 5) proving the mechanistic basis for the higher activity of the -181G promoter.

CREB is a ubiquitously expressed transcription factor belonging to the leucine zipper class of transcription factors with well-known roles in cell proliferation, differentiation, survival and known to be activated by various extracellular stimuli. CREB was initially identified from studies on the cAMP dependent activation of somatostatin gene promoter where it was found to bind an 8 bp cAMP-responsive element (CRE), 5’-TGACGTCA-3’.^33^ Several kinases including protein kinase A (PKA), protein kinase B (AKT), protein kinase C (PKC), MAP kinase activated protein kinase 2 (MAPKAP-2), Ca2+/calmodulin dependent protein kinases II and IV have been shown to activate CREB by phosphorylating the Ser-133 residue which in turn recruits the CREB binding protein (CBP) to activate gene expression.^34^ Targets of CREB include many metabolic enzymes (e.g., Hydroxymethylglutaryl-CoA [HMG CoA] synthase, phosphoenolpyruvate carboxykinase [PEPCK], glucose 6-phosphatase [G6Pase]), neurotransmitters (e.g., Glucagon, tyrosine hydroxylase, chromogranins A and B, enkephalin, vasopressin), transcription factors (e.g., NF-IL-6, Epidermal growth factor-1, STAT-3, c-Fos, PGC-1), growth factors (e.g., Insulin, TNF-α, brain derived neutrophic factor [BDNF], cardiotrophin), cell cycle and survival related proteins (e.g., Retinoblastoma, cyclin A, cyclin D1, Bcl-2), immune regulatory proteins (e.g., Interleukin-2, Interleukin-6, Cox-2) and structural proteins (e.g., fibronectin, αA-crystallin).^34^

Although CREB is known to play an important role in neuropsychiatric disorders, its role in the development of cardiovascular diseases has been recently recognized.^35^ Over-expression of cardiomyocyte-specific dominant negative CREB in transgenic mice under the influence of α-myosin heavy chain gene promoter reduced the cardiac contractility in response to isoproterenol stimulation mirroring idiopathic dilated cardiomyopathy.^36^ Phospho-CREB levels were also found to be elevated in cerebral arteries of hypertensive rats.^37^ Phospho-CREB levels also correlated with the proliferative response resulting in arteriolar injury in Ang II induced hypertension and endothelial injury.^38^ Involvement of CREB in Ang II induced IL-6 expression in vascular smooth muscle cells was attributed to a crucial CRE site. Ang II activated several kinases via AT1-receptor to result in CREB phosphorylation suggesting that CREB may have a role to play in the vascular remodeling associated with cardiac hypertrophy, heart failure and atherosclerosis.^39^

Previous reports have indicated that the *MMP7* promoter, like most MMP promoters, harbors binding sites for activator protein (AP-1) and polyoma enhancer A binding protein-3 (PEA-3) which in turn interact with the Fos and Jun family and Ets family of transcription factors to activate gene transcription.^40,40^ *MMP7* promoter has also been shown to harbor TGF inhibitory elements (TIEs) which are known to bind the SMAD family of transcription factors.^42^Wnt signaling is also known to regulate *MMP7* by binding β-catenin and its partners Tcf/Lef-1.^43^ In the context of the *MMP7* A-181G polymorphism, the higher promoter activity of *MMP7*-181G allele was attributed to the generation of a putative binding site (NGAAN) for a heat-shock transcription factor binding site in case of the -181G allele.^11^ Another study investigating the role of *MMP7* promoter polymorphisms in idiopathic pulmonary fibrosis, wherein the AA genotype was associated with higher plasma levels, reported the interaction of forkhead box A2 (FOXA2) to the -181A allele of the *MMP7* promoter using transient transfection assays and DNA-protein binding assays.^12^ In our earlier study on the association of gastric cancer risk with *MMP7* A-181G polymorphism we had also showed that in the gastric adenocarcinoma cell line, AGS, the higher promoter activity of *MMP7* -181G allele could be attributed to increased binding affinity of CREB to the *MMP7*-181G promoter.^5^ Thus, the current study by virtue of establishing the CREB-mediated activation of *MMP7*-181G promoter in cardiomyoblast and neuronal cells as well sheds light on an important role for CREB in the transcriptional regulation of *MMP7*, especially in cardiovascular diseases.

Since stress elevates catecholamine levels through the hypothalamic-pituitary-adrenal axis^44^ and hypertension is characterized by elevated levels of vasoconstrictive agonists such as catecholamines, we checked if epinephrine, a catecholamine and known activator of CREB exhibited any allele specific effects with respect to the *MMP7*-181A/G promoters. Indeed, the *MMP7*-181G construct displayed a dose-dependent increase in promoter activity in response to increasing doses of epinephrine. The increase in phospho-CREB levels in response to epinephrine treatment and the failure of epinephrine to augment the promoter activity in KCREB co-transfected cells establishes the concomitant role of CREB in epinephrine-mediated elevation in *MMP7*-181G promoter activity (Fig. 6). In the context of chromatin as well, ChIP experiments demonstrated higher promoter occupancy of both CREB and phospho-CREB in case of *MMP7*-181G promoter in the epinephrine treated condition (Fig. 6). Thus, hypertensive carriers of the -181G allele may have even higher expression of *MMP7* gene due to abundance of vasoconstrictive agonists like catecholamines. Interestingly, isoproterenol stimulation was found to activate MMP7 expression at mRNA and protein level in gastric cancer cells by inducing AP-1 and STAT3.^45^ Further, increased expression of MMP7 in gastric cancer tissue was observed at the sites where β2-adrenergic receptor was overexpressed emphasizing the role of epinephrine in activation of *MMP7*-181G promoter in hypertension and other stress induced cardiac complications.

Similarly, hypoxia, a condition resulting in diminished oxygen supply to the tissues is often a pathophysiological occurrence in cardiovascular diseases. HIF-1α, HIF-2α are the most commonly encountered mediators of hypoxia.^46^ HIF-1α activity is regulated at post-translational level wherein under normoxic conditions it undergoes proteasomal degradation while under hypoxic conditions the protein is stabilized and translocated to the nucleus allowing it to bind to the hypoxia responsive elements and activating gene transcription.^47,48^ Hypoxia, in fact, is a common pathophysiological feature in CVDs including atherosclerosis and heart failure. Normotensive Sprague-Dawley rats developed sustained arterial hypertension when subjected to hypobaric hypoxia.^49^ In SHR model of hypertension, the brainstem was found to be hypoxic which lead to increased release of ATP and lactate resulting in elevated sympathetic activity and a corresponding increase in arterial blood pressure.^50^ Chronic hypoxia is often found to be associated with increased sympathetic activity and systemic arterial hypertension. In a study subjecting Danish lowlanders at sea level to an altitude of 5260 m for 9 weeks, the mean blood pressure was found to be 28% higher with up to a 41% increase in DBP accompanied by 3.7-and 2.4-fold higher plasma arterial norepinephrine and epinephrine levels.^51^ HIF-1α signaling has been identified to play important roles in regulation of inflammation (e.g., in macrophage activation) which in turn contributes to the tissue remodeling processes thereby influencing the severity of CVDs.^52^

Although the hypoxic response is primarily mediated by HIF-1α, reports have also shown the involvement of NF-kB and CREB transcription factors. The DNA binding domain of CREB harbors two Cys residues (at positions 300 and 310) which were found to be crucial for the binding of CREB to the CRE site. Mutational analysis revealed that these residues may also act as oxygen sensors resulting in increased expression of CREB-mediated genes in response to hypoxia.^53^ Electrophoretic mobility shift experiments revealed that CREB and ATF-1 were capable of binding to the HIF-1 DNA recognition site.^54^ CREB was further shown to be up-regulated along with ATF-1 and other CREB-responsive genes in response to hypoxia in the lung tissue of mice *in vivo*.^55^ In PC12 cells, hypoxia resulted in Ser-133 phosphorylation of CREB which in turn activated tyrosine hydroxylase. Interestingly, the phosphorylation of CREB observed in response to hypoxia was maximal with a 3-6 hrs exposure and continued to persist for at least 24 hours.^56^ PKA mediated CREB activation was also observed in the ischemic heart.^57^

In line with these reports, hypoxia significantly enhanced the promoter activity of *MMP7*-181G construct under basal as well as CREB co-transfected conditions when compared to cells maintained in normoxia. However, no significant increase in response to hypoxia was observed when the cells were co-transfected with the dominant negative KCREB construct suggesting that the increased promoter activity of *MMP7*-181G construct under hypoxia is modulated by CREB. ChIP experiments revealing increased promoter occupancy of CREB under hypoxic condition in case of the *MMP7*-181G promoter and elevated levels of CREB and phospho-CREB in response to hypoxia further corroborated our findings to confirm the role of CREB in mediating hypoxia induced activation of *MMP7*-181G promoter (Fig. 7). Corroborating our findings, primary human monocyte-derived macrophages subjected to 16 hrs of hypoxia displayed elevated mRNA levels of MMP7 apart from that of VEGF and GLUT-1. Hypoxia also caused an increase in the promoter activity of MMP-7 and GLUT-1 as shown through reporter assays in RAW 264.7 macrophages indicating an activation of gene expression at transcriptional level. MMP7 mRNA and protein levels were shown to be up-regulated in hypoxic macrophages in vitro and in vivo in hypoxic areas of breast tumors.^58^ Thus, the *MMP7* A-181G polymorphism has a functional role in governing gene expression under not only basal condition but also in response to pathophysiological stimuli.

### Genotype-phenotype correlations of *MMP7* A-181G polymorphism

We observed that expression of MMP7 under the influence of *MMP7*-181G promoter was significantly higher than that of *MMP7*-181A promoter suggesting that in the genomic context, the higher promoter activity of the *MMP7* A-181G SNP translated to higher protein levels *in vitro* (Figs. 8 A-E). Plasma MMP7 levels, though modest, were significantly higher in individuals of A/G genotype when compared to wild-type A/A individuals (Fig. 8F) while the plasma MMP7 levels in individuals of variant G/G genotype although higher when compared to wild-type A/A individuals was not statistically significant. Since the heterozygous genotype showed a strong association with hypertension risk in our study populations as well as significantly higher plasma MMP7 levels, we transfected diploid combinations of -181A/-181G constructs mimicking the homozygous and heterozygous combinations *in cella* to study the effect of heterozygous genotype on *MMP7* transcription. Interestingly, the -181A/G heterozygous combination also yielded significantly higher promoter activity than the -181A wild-type indicating that *MMP7* gene expression is likely to be enhanced in the heterozygous individuals as well (Figs. 8G and 8H). Several reports have suggested that altered expression of MMPs in different individuals could result due to the polymorphisms in the regulatory regions (promoter) of MMP genes in these individuals.^2^ Contrary to our finding, the plasma MMP7 levels were higher in individuals of -181AA genotype in idiopathic pulmonary fibrosis (IPF); however, these observations were limited to IPF patients and not in healthy control suggesting that this may be a specific pathological response.^12^

Of note, in a preliminary study, hypercholesterolemic patients with CAD possessing the -181G allele presented with smaller reference luminal diameters before percutaneous transluminal coronary angioplasty than patients with the wild-type allele suggesting a functional role for the *MMP7* A-181G polymorphism in the matrix remodeling associated with CAD.^11^ We also probed for association of the *MMP7* A-181G polymorphism with other physiological/biochemical parameters like total cholesterol, triglycerides, LDL-and HDL-cholesterol and blood glucose levels that may act as co-morbidities to cardiovascular disease; however, no significant correlations were observed. Nonetheless, the significantly higher diastolic blood pressure and mean arterial pressure observed in individuals of A/G genotype (Fig. 9) corroborated with the results from the logistic regression analysis for predicting hypertension risk suggesting plausible contribution for the *MMP7* A-181G polymorphism to hypertension.

In summary, we genotyped a common naturally-occurring polymorphism (A-181G) in the upstream regulatory region of *MMP7* gene in two geographically distinct Indian populations comprising of ∼2500 hypertensive and normotensive subjects using a systematic case-control approach. The *MMP7* A-181G polymorphism showed strong association with increased hypertension risk in our study populations. The -181G allele containing promoter displayed higher promoter activity than the -181A promoter in basal as well as under pathophysiological stimuli (hypoxia, catecholamine excess) due to preferential binding with CREB transcription factor which translated to increased MMP7 levels in vitro (Fig. 10). The risk genotype (-181A/G) also had a correlative association with increased plasma MMP7 levels and blood pressure suggesting that this functional regulatory polymorphism may contribute to cardiovascular risk.

**Figure 10.**
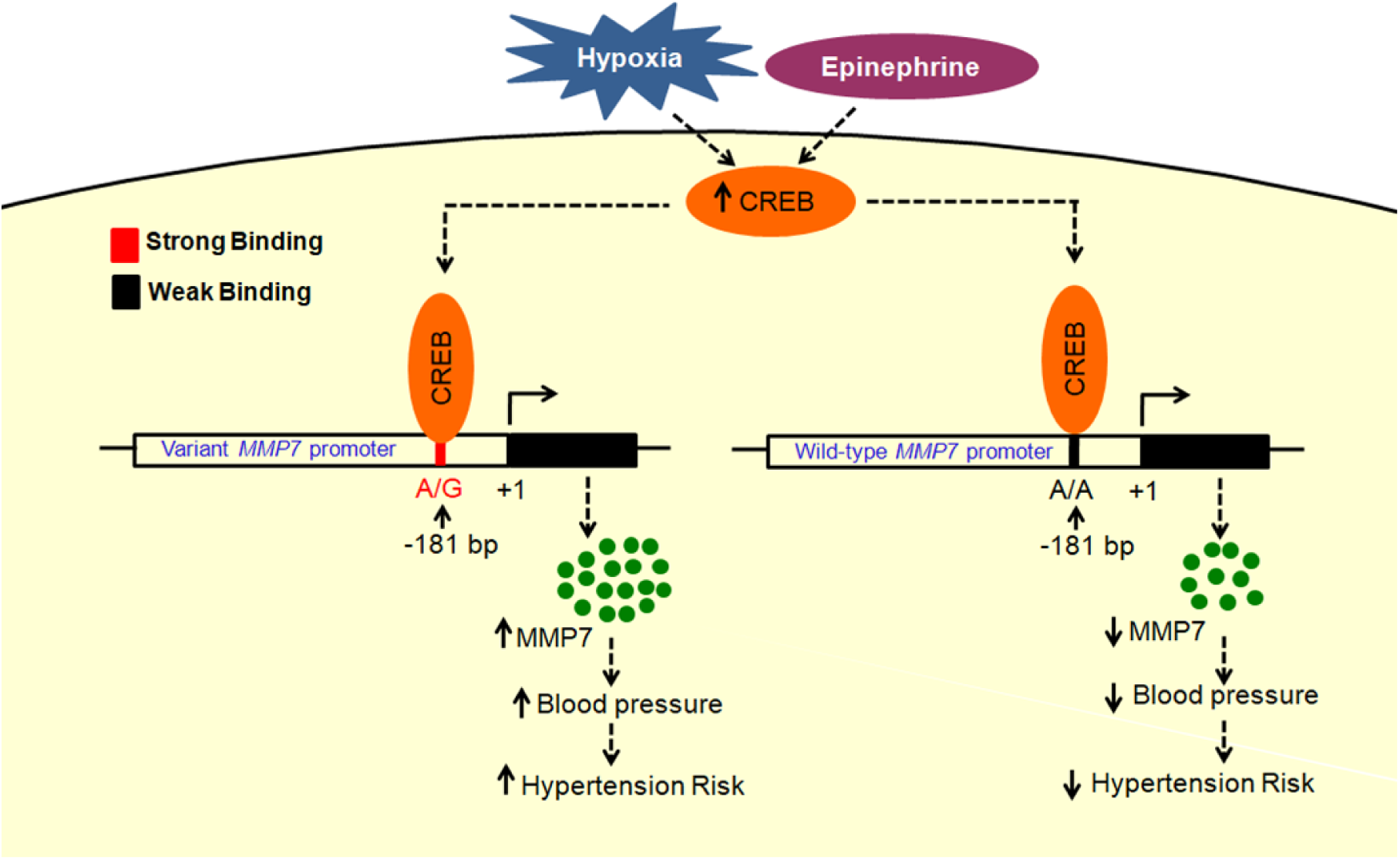
A schematic illustration of the plausible role and mechanistic basis behind the role of *MMP7* -181 A/G promoter variant in contributing to hypertension risk. The *MMP7* -181A/G genotype was found to be associated with increased hypertension risk and elevated blood pressure in the study population with the *MMP7*-181G promoter showing higher promoter activity consistently among different cell lines due to transcriptional activation by CREB. Pathophysiological conditions (viz. catecholamine excess and hypoxia) further increase the promoter activity of *MMP7* -181G construct in an allele-specific manner by activating CREB and inducing its phosphorylation. The higher promoter activity of *MMP7* -181G construct corroborates with the elevated levels of plasma MMP7 observed in -181A/G individuals which, in turn, positively correlates with blood pressure suggesting that individuals heterozygous for the *MMP7* -181A/G genotype may be at a higher risk of hypertension and its complications.

## Perspectives

Uncontrolled proteolytic processes modulated by MMPs resulting in cardiac remodeling plays a major role in the pathological mechanisms involved in hypertension and the resultant cardiovascular complications. MMP7, a potent metalloproteinase with a wide range of ECM and non-ECM substrates majorly implicated in several cancers, atherosclerosis etc. is, therefore, a logical regulator of cardiovascular mortality. This study sheds light on the association of a highly frequent tag SNP in the gene expression controlling promoter region of *MMP7* (A-181G; rs11568818) with increased blood pressure and higher risk for hypertension in Indian population; this work also provides a plausible transcription regulatory mechanism behind the observed elevated levels of MMP7 in the carriers of the variation. Thus, the study paves way towards a better understanding of the role of inter-individual variations and functional regulatory polymorphisms in conferring disease risk, which may be used for developing preventive strategies in individuals likely to be predisposed to hypertension and the related cardiovascular complications.

## Acknowledgments

We acknowledge all the volunteers who participated in this study. We thank Dr. David Ginty (Howard Hughes Medical Institute) for providing the CREB expression plasmid (VP16-CREB+bZIP) and Dr. Richard H. Goodman (Vollum Institute, Oregon Health Sciences University, Portland, OR) for the CREB dominant negative (K-CREB) plasmid. L. S. acknowledges the clinical research team at MMM, Chennai, for helping with the sample collection and Mr. Abrar Ali Khan, Mr. Vikas Arige and Ms. Amrita Anand for their support during the study.

## Novelty and Significance

### What is new?

- This is the first study to the best of our knowledge demonstrating the association of *MMP7* A-181G (rs11568818) polymorphism with hypertension risk.
- This study provides evidence for the role of CREB in the enhanced transcriptional activity of *MMP7* -181G promoter translating to higher protein levels in cardiomyocytes.

### What is relevant?

The study identifies a highly prevalent common tag SNP playing a crucial role in conferring hypertension risk in Indian populations which may be used for developing preventive strategies in individuals likely to be predisposed to hypertension and its complications.

### Summary

We identified a common naturally-occurring polymorphism (A-181G) in the upstream regulatory region of *MMP7* gene showing strong association with increased hypertension risk in two geographically distinct Indian populations comprising of ∼2500 hypertensive and normotensive subjects using a systematic case-control approach. The -181G allele containing promoter displayed higher promoter activity than the -181A promoter in basal as well as under pathophysiological stimuli (hypoxia, catecholamine excess) due to preferential binding with CREB transcription factor which translated to increased MMP7 levels *in vitro*. The risk genotype (-181A/G) also had a correlative association with increased plasma MMP7 levels and blood pressure suggesting that this functional regulatory polymorphism may act as predictor of cardiovascular risk.

### Online Data Supplement

## DETAILED METHODS

### Human Subjects

A total of 2494 unrelated volunteers were enrolled in this case-control study. The primary study population comprised of 862 hypertensive cases (SBP ≥ 140 mm Hg or DBP ≥ 90 mm Hg or a history of hypertension or antihypertensive treatment but no history of cancer or kidney disease) and 655 normotensive controls (with no history of hypertension, diabetes, cancer or kidney disease) from an urban Chennai population recruited at the Madras Medical Mission (MMM), Chennai. The replication study population consisted of 519 hypertensive and 458 normotensive volunteers from Chandigarh recruited at the Postgraduate Institute of Medical Education and Research (PGIMER), Chandigarh. Informed written consent was obtained from each subject and the study was approved by the institutional ethics committee of Indian Institute of Technology Madras. Demographic details (age, gender), physical measurements (height, weight, and body mass index [BMI]), physiological parameters (systolic blood pressure [SBP], diastolic blood pressure [DBP], mean arterial pressure [MAP], heart rate [HR]), biochemical parameters (hemoglobin, sodium, potassium, urea, creatinine, blood sugar, total cholesterol [TC], triglycerides [TGL], low density lipoproteins [LDL] and high density lipoproteins [HDL]) wherever collected from the subjects have been listed in Supplementary Tables 1 and 2). The medical history of the volunteers (current medication, family history of cardiovascular or renal disease states) was recorded at the time of recruitment. Blood samples were collected in EDTA coated tubes by venipuncture for genomic DNA isolation and plasma samples were collected, aliquoted and stored at -80°C for assaying various biochemical parameters. The average age of the cases and controls was ∼44 and ∼42 years in the Chennai study population and ∼49 and ∼59 years in the Chandigarh population respectively. The BMI, blood pressure, sodium levels and triglyceride levels were significantly higher in hypertensive cases than normotensive controls across both the populations.

### Genotyping of *MMP7* -181A/G polymorphism in Indian populations

Genomic DNA was isolated from heparin/EDTA-anticoagulated blood samples using Flexigene DNA kit (Qiagen, USA) according to manufacturer’s protocol. The region of *MMP7* spanning -302 bp to -153bp (NCBI accession number: NM_002423.4; numbered upstream (-) with respect to the cap site) was amplified using the primers: forward, h*MMP7*pro-181FP (5’-TGGTACCATAATGTCCTGAATG-3’) and reverse, h*MMP7*pro-181RP (5’-TCGTTATTGGCAGGAAGCACACAAGAATT-3’). The polymerase chain reaction was performed in Eppendorf Mastercycler. The 150 bp region was amplified in a 10 μl reaction mixture containing 1 mM concentrations of forward and reverse primers, 1.5 mM MgCl_2_, 0.4 mM of each dNTP (New England Biolabs, USA), 20 ng of the template and 0.4 units of the Phusion™ high fidelity DNA polymerase (New England Biolabs, USA). The reaction conditions included an initial denaturation of 2 min at 98°C followed by 35 cycles of denaturation at 98°C for 30s, annealing at 56.5°C for 30s and extension at 72°C for 30s with a final extension step at 72°C for 10 min. The PCR products were digested with EcoRI enzyme and the digested products were electrophoresed in a 2.5% agarose gel. The genotypes were inferred by observed the digested band pattern. The presence of variant (G allele) creates a restriction site for EcoRI which digests the 150 bp PCR product to yield two fragments of size 120 bp and 30 bp, while the wild-type (A allele) remains undigested. Heterozygous (A/G) genotype results in three bands of sizes 150 bp, 120 bp and 30 bp respectively. About 10% of the PCR products were also purified and sequenced by Sanger sequencing to confirm the inferred genotypes.

### Estimation of biochemical parameters

Biochemical parameters such as random blood sugar, total cholesterol, triglycerides, LDL, HDL, urea, creatinine, hemoglobin, sodium and potassium levels were measured in the plasma using standard biochemical assays. The blood pressure readings were measured by experienced nursing staff as an average of triplicate values recorded in the sitting position using a brachial oscillometric cuff. Plasma MMP7 levels were measured by a commercially available immunoassay kit (R&D systems, USA) following the manufacturer’s protocol.

### Cloning and mutagenesis

The ∼250 bp promoter region of *MMP7* was amplified by PCR using genomic DNA of known homozygotes for *MMP7*-181A/G polymorphism and the following primers: h*MMP7*-230-FP: 5’-CGG**GGTACC**TGGAGTCAATTTATGCAGCAGACAG-3’ and h*MMP7*+22-RP: 5’-CCG**CTCGAG**TTGGACCTATGGTTGATTTGGTG-3’ (restriction sites of KpnI and XhoI inserted at the 5’ end of forward and reverse primers respectively are indicated in boldface). The purified promoter fragments were inserted between KpnI and XhoI sites in the promoterless luciferase reporter vector pGL3-Basic which has the firely luciferase as the reporter gene (New England Biolabs, USA). The generated promoter-reporter constructs consisted of the -230-to +22-bp region of *MMP7*. Accurate cloning of the inserts with the variation was confirmed by DNA sequencing using the primers: RVprimer3 5’-CTAGCAAAATAGGCTGTCCC-3’ and GLprimer2 5’-CTTTATGTTTTTGGCGTCTTCCA-3’.

To generate the *MMP7* promoter haplotype-cDNA constructs, *MMP7* cDNA was subcloned from pcDNA3-GFP-MMP7, a gift from Steven Johnson (Addgene plasmid # 11989) into the *MMP7*-181G and *MMP7*-181A promoter-reporter constructs replacing the luciferase cDNA. The *MMP7*-181G and *MMP7*-181A constructs were digested with NcoI followed by mungbean nuclease treatment to blunt the sticky end and then digested with XbaI to release the 1.6 kb luciferase cDNA. The pcDNA3-GFP-MMP7 plasmid was digested with EcoRI followed by mungbean nuclease treatment to blunt the sticky end and then digested with XbaI to release the 804 bp MMP7 cDNA. The 804 bp MMP7 cDNA was then ligated into the 3.1 kb backbone of *MMP7*-181G and *MMP7*-181A constructs. The clones were confirmed by sequencing using primers h*MMP7*-230-FP: 5’-CGGGGTACCTGGAGTCAATTTATGCAGCAGACAG-3’ and pGL3pro RP: 5’-CCCCCTG AACCTGAAACATA-3’.

All the plasmids were purified on columns using an endotoxin-free plasmid DNA purification kit (Hi Media, India) for transfection experiments.

### Cell culture, transfection and reporter assays

Human neuroblastoma IMR-32, SH-SY5Y, rat cardiomyoblast H9c2 and mouse neuroblastoma N2a cell lines were obtained from the National Center for Cell Sciences, Pune, India. Cell lines were maintained in Dulbecco’s Modified Eagle Medium (DMEM) containing high glucose, L-glutamine and sodium pyruvate (HyClone, USA) supplemented with 10% (v/v) fetal bovine serum (Gibco, USA), penicillin G (100 U/ml) and streptomycin sulfate (100 mg/ml) (Gibco, USA) in humidified incubators at 37°C with 5% CO_2_(Thermo Scientific, USA). Cells were grown up to 70% confluence in 24-well plates and transfections were performed in triplicates with 1.0 μg/well of promoter-reporter plasmid and 300 ng/well of a β-galactosidase expression plasmid (as internal control). Cells were transfected using Targefect F2 transfection reagent (Targeting Systems, USA) following manufacturer’s protocol. The culture media was changed to fresh complete medium after 6 hrs. After 30 hrs of transfection, cells were lysed and luciferase and β-galactosidase assays were performed with the cell lysates as described previously.^1^ Promoter activities were expressed as luciferase/β-galactosidase readings.

Co-transfection experiments were carried out in IMR-32 and H9c2 cells with different doses of CREB expression plasmid (a kind gift from Dr. David Ginty, Howard Hughes Medical Institute, USA)^2^ along with *MMP7*-181G and *MMP7*-181A promoter-reporter constructs. pcDNA3.1(+) was used as balancing plasmid. Similarly co-transfection experiments with different doses of dominant negative KCREB expression plasmid (a kind gift from Dr. Richard H. Goodman, Vollum Institute, Oregon Health Sciences University, USA)^2^ along with *MMP7*-181G and *MMP7*-181A promoter-reporter constructs were carried out in IMR-32 and H9c2 cells. Human CREB dicer substrate siRNA oligos (sense, 5’-CCACCAAUCUGCUUCCUGUUUCUTT-3’ and antisense, 5’-AAAGAAACAGGAAGCAGAUUGGUGGUU-3’) and negative control siRNA oligos (Integrated DNA Technologies, Belgium) were co-transfected in 12-well plates with *MMP7*-181G and *MMP7*-181A promoter-reporter plasmids using Targefect F2 transfection reagents (Targeting Systems, USA) into IMR-32 cells. Similarly, CREB dicer substrate siRNA oligos (sense, 5’-AAAGAAACAGGAAGCAGAUUGGUGGUU-3’ and antisense, 5’-AAGGAAUUUCCCAAGUUGCUGAUACCC-3’) were used for siRNA co-transfection experiments in H9c2 cells. In all co-transfection experiments, cell lysates were assayed for luciferase activity 36 hrs post-transfection. Total protein per individual well was also estimated in the corresponding cell lysates by using Bradford reagent (Bio-Rad, USA) and reporter activities were expressed as luciferase activity/μg of protein.

In another set of experiments, IMR-32 and H9c2 cells transfected with the *MMP7*-181G and *MMP7*-181A promoter-reporter constructs were treated with different concentrations of epinephrine (0.5, 1.0 and 5.0 μM) in serum free media after 24 hrs of transfection. In some experiments, KCREB expression plasmid (1.0 μg/well) was co-transfected in IMR-32 and H9c2 cells prior to treatment with epinephrine (5.0 μM). Cell lysates were assayed for luciferase activity after 6 hrs of treatment with epinephrine. The promoter activities were normalized with total protein.

For hypoxia experiments, *MMP7*-181G and *MMP7*-181A promoter-reporter constructs were transfected into IMR-32 and H9c2 cells alone or co-transfected with CREB or KCREB expression plasmid and after 24 hrs of transfection, culture media was changed to serum free media and the cells to be maintained under hypoxic condition were transferred to a desiccator cabinet (Bel-Art, USA) which was flushed with argon gas for 3 min and sealed. The cells under this hypoxic condition were placed in an incubator at 37°C. In parallel, for normoxic condition, 24 hrs after transfection, culture media was replaced with serum free media and the cells were transferred to a regular CO_2_ incubator. Luciferase and Bradford assays were performed after 12 hrs of hypoxia.

In certain experiments, *MMP7* promoter-reporter plasmids were transfected in diploid combinations to imitate the homozygous and heterozygous conditions *in cella*. IMR32 and H9c2 cells were transfected with 1 μg each of *MMP7*-181G or *MMP7*-181A or 500 ng each of *MMP7*-181G and *MMP7*-181A constructs along with β-galactosidase expression plasmid. Luciferase and β-galactosidase assays were performed after 24 hrs of transfection. Promoter activities were expressed as luciferase/β-galactosidase readings.

In another set of experiments, -181G-MMP7cDNA and -181A-MMP7cDNA constructs were transfected into N2a and H9c2 cells and the MMP7 levels in the protein lysates were estimated after 48 hrs of transfection by western blotting to assess the differences in transcription of *MMP7* under the influence of *MMP7*-181A/G SNP.

### Western blotting

Over-expression or down-regulation of CREB, phospho-CREB and HIF-1α after transfection experiments/epinephrine treatment/hypoxia was determined by western blotting. The cells were lysed in a buffer containing 50 mM Tris-HCl (pH 7.4), 150 mM NaCl, 1% Triton X-100, 1% sodiumdeoxycholate, 1 mM EDTA, 0.1% SDS, 1 mM PMSF, protease inhibitor mixture (Sigma) and phosphatase inhibitors (1 mM sodium orthovanadate, 1 mM sodium pyrophosphate and 50 mM sodium fluoride) followed by sonication. The protein concentrations were estimated by Bradford assay (Bio-Rad, USA). Also, *MMP7* promoter–cDNA constructs were transfected into N2a and H9c2 cells and the total protein was isolated after 48 hrs of transfection and subjected to western blotting. Equal amount of protein samples (50 μg) per condition were subjected to SDS-PAGE on a 10% resolving gel and transferred to nitrocellulose membrane (Pall Life Sciences, Mexico). The membranes were blocked with 3% BSA or 5% non-fat milk for 1 hr at room temperature followed by incubation with specific primary antibody [CREB (Cell Signaling Technology, #9197) at 1:1500 dilution, phospho-CREB (Cell Signaling Technology, #9198) at 1:1500 dilution, HIF-1α (Cell Signaling Technology, #14179) at 1:1500 dilution, MMP7 (Abcam, ab5706) at 1:1000 dilution, Vinculin (Sigma, V9131) at 1:10000 dilution overnight at 4°C. After washing with 1xTBST, the membrane was incubated with HRP-conjugated secondary antibody specific for either rabbit (Jackson Immunoresearch Laboratories Inc., 111-035-003 at a dilution of 1:5000 for CREB, phospho-CREB, HIF-1α and MMP7) or mouse (Jackson Immunoresearch Laboratories Inc. # 115-035-003 at 1:5000 dilution for Vinculin) for 1 hr. The detection was carried out using an enhanced chemiluminescence kit (Bio-Rad, USA) and the luminescence signal was captured using Chemidoc XRS+Chemiluminescence Detection system (Bio-Rad, USA). The intensities of signals were quantified using Image Lab software (Bio-Rad, USA).

### Chromatin immunoprecipitation (ChIP) assays

ChIP assays were carried out as described earlier^3^. Briefly, N2a and H9c2 cells were transfected with *MMP7*-181G and *MMP7*-181A promoter-reporter constructs and cross-linkedin 1% formaldehyde in PBS after 24 hrs of transfection. Chromatin was isolated and sheared by sonication, followed by pre-clearing with Rec protein G-Sepharose 4B beads (Invitrogen, USA). Pre-cleared samples were then immunoprecipitated by incubation with 5 μg each of CREB antibody. Non-immune Rabbit IgG (I5006, Sigma) was used as a negative control. The immunoprecipitated chromatin was captured with Rec protein G-Sepharose 4B beads, eluted, reverse cross-linked, digested with Rnase and proteinase K and purified using HiPurA^™^ PCR product purification kit (HiMedia, India).

Following purification, PCR was carried out to amplify the DNA sequence encompassing the *MMP7*-181A/G site of polymorphism using the following primers (forward [h*MMP7*ChIP-230FP], 5’-TGGAGTCAATTTATGCAGCAGACAG-3’ and reverse [h*MMP7*ChIP+22RP], 5’-TTGGACCTATGGTTGATTTGGTG-3’) followed by agarose gel electrophoresis of the PCR products. ChIP quantitative PCR was performed using DyNAmoColorFlash SYBR Green kit (Thermo Scientific, USA) using the same primers. The amounts of DNA immunoprecipitated by CREB due to binding to the -181G and -181A alleles were quantified by the fold enrichment method relative to IgG signal.

In another set of experiments, N2a cells transfected with *MMP7*-181G and *MMP7*-181G promoter-reporter constructs were subjected to treatment with 5.0 μM epinephrine or hypoxia treatment followed by isolation of chromatin for ChIP assays. Immunoprecipitation was carried out with 5 μg each of CREB and phospho-CREB antibodies (IgG was used as negative control) followed by qPCR of purified immunoprecipitated DNA to quantify fold enrichment relative to IgG.

### Building the homology model of CREB1

Sequence of hCREB1 (UniProtID: P16220) was taken from UniProt (http://www.uniprot.org). Disorder regions and protein binding regions were predicted through DISOPRED3^4^ server and DNA-binding residues were predicted using DP_Bind^5^ and PredictProtein^6^ server. Then final homology models of human CREB1 (residue 280-341) monomer and dimer (with Mg^2+^ ion), were built through the MODELLER v11.9^7^, using the template PDB-id. 1DH3 (285-339, *Mus musculus*, mCREB1, resolution: 2.8 Å). hCREB1 and mCREB1 showed 99.7% sequence conservation. Both have 341 amino acid long CREB1, and out of this 340 residues are conserved except residue at 6^th^ position. Two hundred conformations were generated and the best model with highest negative DOPE_score was selected and further refined and minimized by using OPLS3 force field^8^ through protein-preparation-wizard panel in Maestro v11.2.014^9^. Model’s quality assessment was done via Ramachandran plot^10^, ProSA_Zscore^11^, ERRAT^12^, Dope_Score^7^ and secondary structure analysis.

### Building of *MMP7* promoter DNA models

Models of *MMP7* promoter DNA of 45bp (±22bp flanking -181bp position) for *MMP7*-wild-type with -181A allele and *MMP7*-mutant with -181G allele were built using the tools 3DDART^13^ and SCFBio (http://www.scfbio-iitd.res.in/software/drugdesign/). Additionally, the DNA models were further refined through Schrodinger of Maestro v11.2.014 by using force field OPLS3^8^. Furthermore, the 100 ns molecular dynamics (MD) simulation was performed (DESMOND tool of Schrodinger Maestro), to check the stability of DNA in aqueous solution.

### Docking of CREB1 to *MMP7* promoter DNA

Protein–DNA docking was performed with HDOCK^14^ and HADDOCK^15^ tools. In protein-DNA docking with interface information, HDOCK server ranked within the top performed group in CAPRI experiment 2017. HDOCK server utilized hybrid docking algorithm of template-based modeling and free docking. It used hierarchical FFT-based global docking protocol with an improved shape-based pair-wise scoring function to globally sample putative binding modes. HADDOCK generated maximum 200 protein-DNA docking complexes (water-refined models) and clustered them according to their HADDOCK score, more negative score being the more reliable complex. The weighted sum of intermolecular electrostatic (Elec), van der Waals (vdW), desolvation (Dsolv) and ambiguous interaction restraints (AIR) energies and buried surface area (BSA) Term, in total formed HADDOCK score. In HADDOCK server, docking was performed in three successive stages; first, rigid-body docking was performed which generated all possible combination of protein-DNA complex of which top 20% (based on HADDOCK scores) of the complexes were selected and further refined in semi-flexible refinement stage and finally all filtered complexes were refined in explicit solvent. Z-score of cluster indicates the standard deviation from the average (the more negative the better).

### Molecular dynamics simulation of CREB1-MMP7 complexes

Molecular dynamics (MD) simulation for all CREB-*MMP7* (protein-DNA) complexes/models was carried out using Desmond^16^ tool in Maestro. System Builder panel was used to build molecular systems. Inbuilt OPLS3 force field was used to assign all parameters. System was solvated with TIP3P^17^ water molecules using orthorhombic box with distance of 10Å from all sides of protein complex. Orthorhombic box volume was minimized by reorientation of the complex/model system. Appropriate number of counter Na^+^/Cl^−^ions was added to electrically neutralized system. The Steepest descent algorithm with 2000 iterations and convergence threshold of 1 kcal/mol/Å was used to minimize all prepared systems, then further equilibration was performed by using default algorithm which include two stage of minimization and 4 stage of MD. Finally, 100 ns of molecular dynamics simulation was performed using NPT ensemble with temperature and pressure coupling of 300 K and 1 atm respectively. Coordinates and energy were recorded every 10 ps to yield 10000 frames.

MM-GBSA (Molecular Mechanics, the Generalized Born model and Solvent Accessibility) calculation was performed through Schrodinger by using thermal_mmgbsa.py script with step size of 100. A total 100 frames out of 10000 frames were selected for hydrogen bond energy and electrostatic energy calculation. The inbuilt prime MM-GBSA module with OPLS3 force field and generalized Born implicit solvent model in Schrodinger was used to calculate average binding free energy (ΔG).

### Data presentation and statistical analysis

*MMP7* upstream regulatory variants data was collected from Ensemble database. Linkage disequilibrium (LD) analysis was carried out using Haploview 4.2^18^. Phenotypic parameters of the study population were expressed as mean ± S.E. Statistical Package for Social Sciences (SPSS) Version 21.0 was used to carry out genotype-phenotype association studies where the statistical significance of the associations were tested using one-way ANOVA with post hoc tests or Levene’s test for equality of variances followed by two-tailed t-test, as appropriate. Transcription factor binding site prediction was carried out using ConSite^19^, MatInspector^20^ and P-Match^21^. Promoter-reporter transient transfections and co-transfections were carried out at least three times and results were expressed as mean ± S.E. of triplicates from representative experiments. Statistical significance was calculated by Student’s t-test or one-way ANOVA with Bonferroni’s multiple comparisons post-test or two-way ANOVA, as applicable, using Prism 5 program (GraphPad Software Inc., USA). All representative graphs were generated using GraphPad Prism 5 software.

## SUPPLEMENTARY RESULTS

### Genotyping of *MMP7* -181 SNP in Indian populations

Genotyping for the *MMP7* A-181G SNP was carried out by PCR-restriction fragment length polymorphism (RFLP) method using EcoRI restriction enzyme. Undigested amplicons of 150 bp indicated -181A allele and EcoRI restricted fragment sizes of 120 bp and 30 bp represented the -181G allele (Fig. S1). A subset of the samples was subjected to Sanger DNA sequencing (Fig. S1) for confirmation of genotypes. Genotyping results from both these techniques showed 100% concurrence.

### Generation and assessment of the quality of CREB1 model structure

While the complete sequence of hCREB1 consists of 341 amino acids (aa) crystal structure of only the 55 aa stretch of CREB1 protein (285-339 aa, PDB ID: 1DH3, species: *Mus musculus*, identity: 97.7%) is available. Therefore, we set out to build a model structure of the transcription factor (TF) hCREB1 for interaction studies with the MMP7 promoter DNA.

First, we carried out *in-silico* structural profiling of CREB1. Since TFs are known to have long range of intrinsic-disorder zone and undergoes to disorder-to-order transitions upon binding to other partners proteins,^22-24^ we carried out disorderedness analysis on CREB1. DISOPRED3^4^ was utilized to predict the disordered regions in CREB1. In DISOPRED score, the region with confidence score > 0.5 was considered as disordered one. The amino acid residues from 1 to 160 were highly disordered (confidence score > 0.8), residue 211 to 280 were moderately disordered (confidence score: 0.6-0.8), and residue 161 to 210, 281 to 350 were structured (confidence score < 0.5). This finding is in line with the non-availability of crystal structures for CREB1.

Next, we analyzed the CREB1 sequence to predict DNA binding residues by the DP-Bind^5^ and PredictProtein^6^ tools. The results of DP_Bind and PredictProtein matched well with the binding reported in the PDB ID: 1DH3. It was found that the regions of the protein, which mostly interacted with DNA, had stable and fixed secondary structure^22^. From the results of disorderedness prediction and from the DNA-binding, it was clear that only 280-341 region had both properties (viz. fixed secondary structure and DNA binding affinity). Therefore, we generated CREB1 homology model for only this region (280-341).

Homology model of human CREB1 (hCREB1) residues 280-341, was generated through MODELLER v11.910. The final model had 86.8% residues in the most favored regions in the Ramachandran plot while in the *Mus musculus* CREB1 crystal structure template 1DH3, 73.6% residues occurred in most favored region and none of the residues were found in the disallowed region in both the structures. The Prosa Z-score for the modeled CREB1 is −1.87 and −0.97 for CREB template (1DH3). The overall quality factor in ERRAT was 97.8 for the modeled CREB1 and 93.7 for the CREB template 1DH3. The Dope_score and secondary structure predictions for the modeled structure also showed a trend similar to that of the crystal structure (1DH3). The C-alpha RMSD analysis (data generated through molecular dynamics simulation) between HD region of model-vs.-1DH3 was 1.1 Å, indicating the robustness and good quality of the model.

### CREB1 and *MMP7*-DNA docking predicted greater binding affinity of the mutant promoter

Focused protein-DNA docking of *MMP7*-wild-type and *MMP7*-mutant (mut) DNA molecules with CREB1 was performed to assess the effect of the A-181G mutation on binding affinity. As shown in the Supplementary Table 4, the CREB1:*MMP7*-mut complex had relatively higher HADDOCK score (-138.280+/-2.7) and larger cluster size (total complex = 80) as compared to the CREB1:*MMP7*-wild-type complex score (-136.280+/-3.6) and cluster size (total complex= 74) suggesting that *MMP7*-mut (-181G) DNA may have higher binding affinity as compared to the *MMP7*-wild-type (-181A) DNA for CREB1. Similar results were also obtained from HDOCK with the best complex having a docking score of -1120.96 and -1110.66, for CREB1:*MMP7*-mut and CREB1:*MMP7*-wild-type, respectively.

Residues forming stable hydrogen bonds with distance up to 3.5Å and more than 30% occurrence throughout 100 ns MD trajectory were considered as hot-spot residues. Based on the MD simulations, residues N293 and R301 were identified as hot-spot residues in the CREB1:*MMP7*-wild-type while in CREB1:*MMP7*-mut R289 was also identified as a hot-spot residue in addition to N293 and R301.

Of note, structural changes of *MMP7*-wild-type and *MMP7*-mut DNA in aqueous system were observed (Figs. S3A and S3B). During MD simulation, the hCREB1(monomer):*MMP7*-promoter complex also displayed high fluctuations. Therefore, CREB1(dimer):*MMP7*-wild-type and CREB1(dimer):*MMP7*-mut complexes were built. From the MD simulations, stable complexes with the lowest RMSD were selected for further analysis (Fig. S3; panels C1, C2 and C3). Both the *MMP7*-wild-type (average RMSD, 6.9 Å) and *MMP7*-mut (average RMSD, 6.4 Å) DNA evolved simultaneously from 2 to ∼7 ns and then became stable till 100ns. Both the complexes follow the same trend in MD analysis although RMSD calculations suggested that *MMP7*-mut was more stable than *MMP7*-wild-type; these results may be attributed to the increased GC content (in the case of the mutant DNA) that improves thermal stability of DNA. Changes in the trajectory of *MMP7*-wild-type (a shift in RMSD from 2.5 Å to 6.4 Å) were observed; however, in case of *MMP7*-mut, the trajectory remained consistent with value of 2.4 Å throughout 100ns. Furthermore, a bimodal *vs.* unimodal distribution was observed in *MMP7*-wild-type *vs MMP7*-mut (Fig. S3C1). Chain-wise analysis of CREB1(dimer):*MMP7* complex revealed that the changes in RMSD of *MMP7*-wild-type occurred mainly due chain A where the RMSD distribution showed a significant shift from 3.0 Å to 8.0 Å (bimodal distribution) (Figs. S3C2 and S3C3). In case of *MMP7*-mut, the RMSD distribution of chain A remained consistent at 3.0 Å throughout the MD simulation and had a unimodal distribution. The Root mean square fluctuation (RMSF) of the C? atom of the entire residue over different time frame was plotted to measure the fluctuation of the interface residues over the simulation time for *MMP7*-wild-type and *MMP7*-mut systems (chain A and B, separately).

We further studied the regions that showed significant structural changes during MD simulations by determining the RMSF difference between *MMP7*-wild-type and *MMP7*-mut (Fig. S3D). The RMSF difference values for chain B is comparatively smooth and consistent which clearly indicates that the interaction between chain B and DNA is stable in both cases. However, in case of CREB1 chain A:*MMP7*-wild-type, the RMSF difference had a huge fluctuation when compared to CREB1 chain A:*MMP7*-mut, validating the outcome of RMSD. The residues that showed large fluctuations in the case of chain A:*MMP7*-wild-type complex ranged from amino acids 285-292, 305-315 and 334-339. Distance analysis between the *MMP7*-181 A/G nucleotide and the “center of mass” of residues 285-302 of chain A of CREB1 revealed that the distance was consistent throughout the trajectory with unimodal distribution (mean average distance is 14.0 Å) in case of *MMP7*-mut; on the other hand, the distance varied after 40ns and it increased to 25 Å trimodal distribution in case of *MMP7*-wild-type (Fig. S3E).

Next, we carried out detailed hydrogen bond analysis in *MMP7*-wild-type and *MMP7*-mut complexes. While the total numbers of hydrogen bonds were similar in both *MMP7*-wild-type and *MMP7*-mut systems up to 20 ns, the number increased in the case of *MMP7*-mut but not in the case of *MMP7*-wild-type between 20 – 35 ns. Interestingly, after 40 ns (till 100 ns), the total number of hydrogen bonds in the case of *MMP7*-mut remained consistent and continuous, while no hydrogen bonds were observed in the case of *MMP7*-wild-type indicating loss of contacts (Fig. 3F). Further, to observe the net thermodynamic changes between *MMP7*-wild-type and *MMP7*-mut systems, the MM-GBSA energy calculation between CREB1 (chain A) and *MMP7* -181 nucleotides A/G was performed. The total electrostatic energy of both systems remained favorable (negative) till 40 ns, however after 40 ns, the energy became positive in the case of *MMP7*-wild-type, indicating displacement of chain A from the -181A nucleotide (Fig. S3G). Taken together, our extensive computational analysis provided the molecular basis for enhanced interactions of the *MMP7* mutant promoter with CREB as compared to the MMP7 wild-type promoter.

## SUPPLEMENTARY TABLES

**Table S1.** Clinical characteristics of the Chennai study population.

| Characteristics | Hypertensive | Control |  |
| --- | --- | --- | --- |
| a) Demographic | Value ± SD (n) | Value ± SD (n) | p value |
| Age (years) | 44±9 (860) | 42±9 (638) | <0.01 |
| Sex (M/F) (%) | 70.7/29.3 (861) | 71.1/28.9 (640) |  |
| Body Mass Index (kg/m<sup>2</sup>) | 25.4±3.6 (840) | 25.9±3.6 (611) | <0.05 |
| b) Physiological | Value ± SEM (n) | Value ± SEM (n) | p value |
| Systolic Blood Pressure (mmHg) | 145.7±0.6 (815) | 122.3±0.5 (628) | <0.0001 |
| Diastolic Blood Pressure (mmHg) | 89.7±0.4 (819) | 77.9±0.3 (627) | <0.0001 |
| Mean Arterial Pressure (mmHg) | 108.2±0.4 (815) | 92.8±0.3 (627) | <0.0001 |
| Heart rate | 76.1±0.3 (772) | 75.7±0.3 (585) | 0.44 |
| c) Biochemical | Value ± SEM (n) | Value ± SEM (n) | p value |
| Sodium (meq/L) | 138.9±0.2 (348) | 139.5±0.1 (368) | <0.05 |
| Potassium (meq/L) | 4.28±0.02 (345) | 4.32±0.02 (364) | 0.15 |
| Urea (mg/dl) | 21.7±0.2 (762) | 21.2±0.2 (562) | 0.21 |
| Creatinine (mg/dl) | 0.84±0.01 (772) | 0.82±0.01 (565) | 0.09 |
| Random Blood Sugar (mg/dl) | 111.1±1.4 (752) | 104.2±1.3 (550) | <0.001 |
| Total Cholesterol (mg/dl) | 176.8±1.4 (743) | 182.2±1.5 (546) | <0.05 |
| Triglycerides (mg/dl) | 156.1±2.9 (743) | 139.0±2.8 (543) | <0.0001 |
| HDL (mg/dl) | 38.1±0.3 (742) | 38.8±0.3 (540) | 0.12 |
| LDL (mg/dl) | 109.3±1.3 (747) | 115.2±1.4 (551) | <0.01 |

**Table S2.** Clinical characteristics of the Chandigarh study population.

| Characteristics | Hypertensive | Control |  |
| --- | --- | --- | --- |
| a) Demographic | Value ± SD (n) | Value ± SD (n) | p value |
| Age (years) | 49±12 (493) | 59±8 (456) | <0.0001 |
| Sex (M/F) (%) | 52.4/47.6 (494) | 70.0/30.0 (457) |  |
| b) Physiological | Value ± SEM (n) | Value ± SEM (n) | p value |
| Systolic Blood Pressure (mmHg) | 147.9±0.6 (492) | 118.8±0.5 (441) | <0.0001 |
| Diastolic Blood Pressure (mmHg) | 95.5±0.4 (508) | 75.3±0.3 (442) | <0.0001 |
| Mean Arterial Pressure (mmHg) | 113.1±0.4 (502) | 89.8±0.3 (440) | <0.0001 |
| c) Biochemical | Value ± SEM (n) | Value ± SEM (n) | p value |
| Sodium (meq/L) | 139.6±0.5 (100) | 142.8±0.9 (58) | <0.01 |
| Potassium (meq/L) | 4.70±0.42 (98) | 5.03±0.64 (58) | 0.65 |
| Urea (mg/dl) | 31.1±0.9 (149) | 33.8±0.9 (61) | 0.10 |
| Creatinine (mg/dl) | 1.26±0.24 (151) | 0.92±0.04 (62) | 0.36 |
| Random Blood Sugar (mg/dl) | 94.2±1.0 (97) | 96.8±1.4 (58) | 0.13 |
| Total Cholesterol (mg/dl) | 190.3±2.0 (454) | 185.5±4.3 (65) | 0.31 |
| Triglycerides (mg/dl) | 163.6±2.7 (456) | 125.7±4.1 (64) | <0.0001 |
| HDL (mg/dl) | 46.5±0.5 (454) | 41.7±1.1 (65) | <0.001 |
| LDL (mg/dl) | 129.8±1.9 (460) | 143.1 ±5.0 (65) | <0.05 |

**Table S3.** SNPs in the 5 kb upstream promoter region of *MMP7*.

| Variant ID | Chromosome position | *MMP7* Promoter Position | Minor Allele Frequency in Overall Population (n=2504) | Minor Allele Frequency in South Asian (SAS) Population (n=489)<sup>25</sup> |
| --- | --- | --- | --- | --- |
| rs11568819 | 11:102530902 | -153 | 0.040 | 0.047 |
| **rs11568818** | **11:102530930** | **-181** | **0.358** | **0.443** |
| rs17098318 | 11:102532127 | -1378 | 0.301 | 0.370 |
| rs17881620 | 11:102532522 | -1773 | 0.255 | 0.328 |
| rs10895308 | 11:102532844 | -2095 | 0.209 | 0.083 |
| rs72982143 | 11:102533418 | -2669 | 0.038 | 0.044 |
| rs2408391 | 11:102533760 | -3011 | 0.238 | 0.270 |
| rs72982145 | 11:102534227 | -3478 | 0.042 | 0.053 |
| rs880197 | 11:102534940 | -4191 | 0.238 | 0.270 |
| rs1943778 | 11:102535234 | -4485 | 0.031 | 0.039 |

**Table S4.**
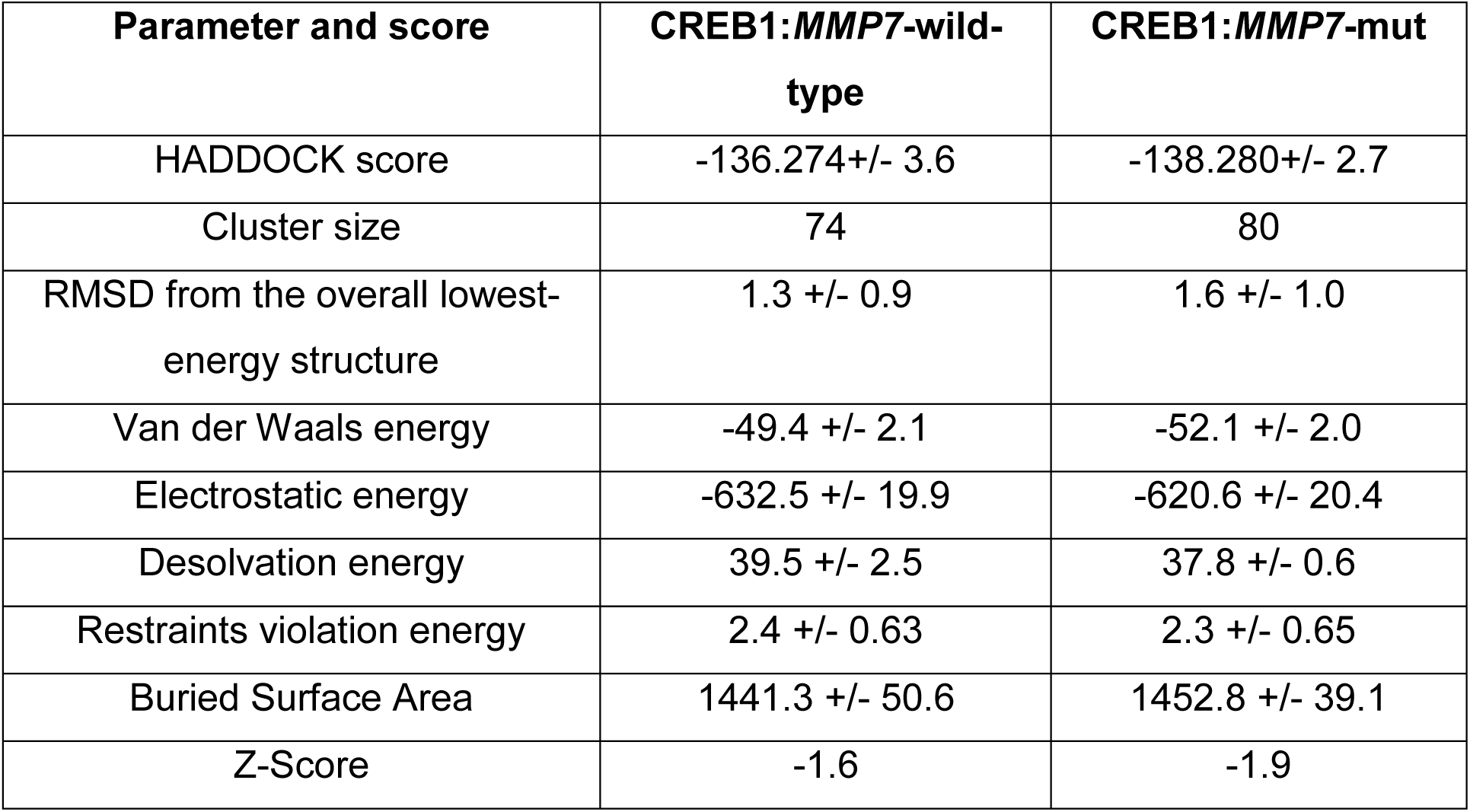
Results of focused protein-DNA docking of CREB1 (monomer) and *MMP7*-promoter DNA through HADDOCK server.

## SUPPLEMENTARY FIGURES

**Figure S1.**
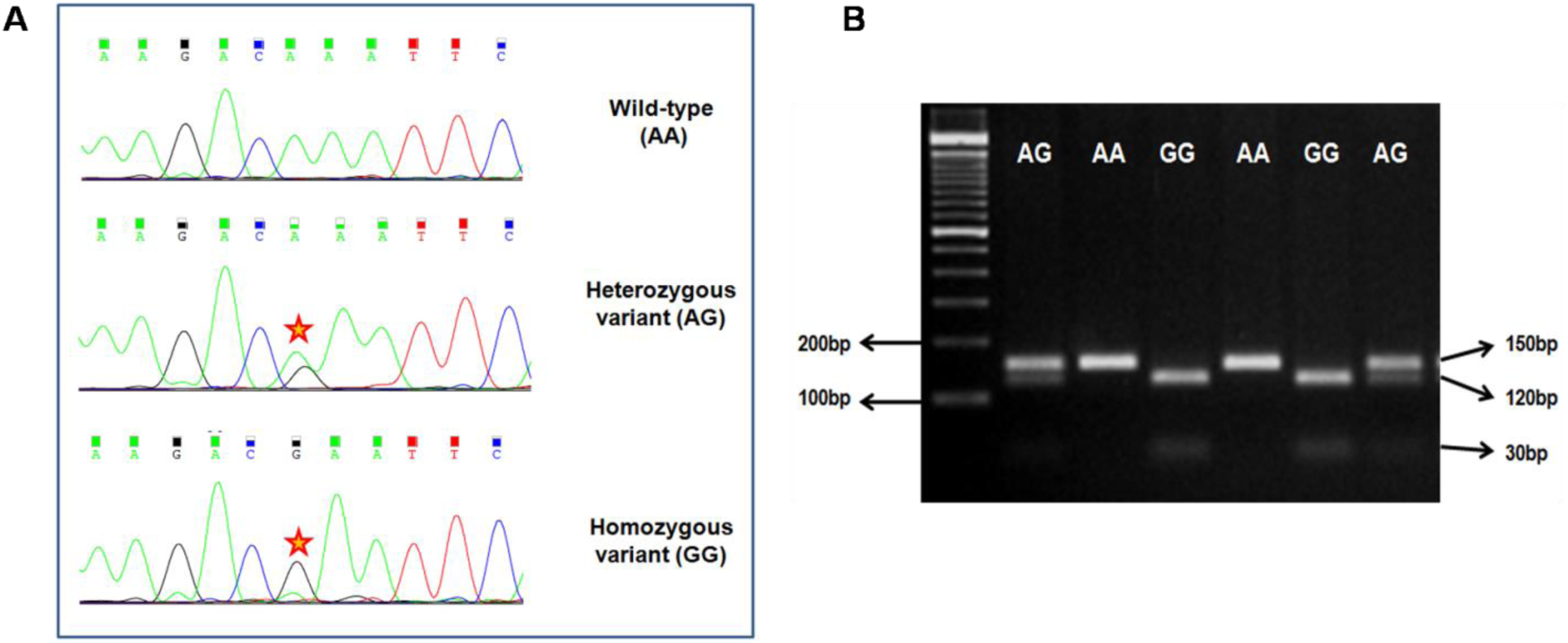
Genotyping of *MMP7*-181A/G SNP. **(A)** Chromatograms representing the AA, AG and GG genotypes of *MMP7*-181A/G SNP (rs11568818). **(B)** RFLP-genotyping of rs11568818 using EcoRI. The 150 bp PCR product flanking the SNP is cleaved in case of G allele but not the A allele. Inferred genotypes are indicated.

**Figure S2.**
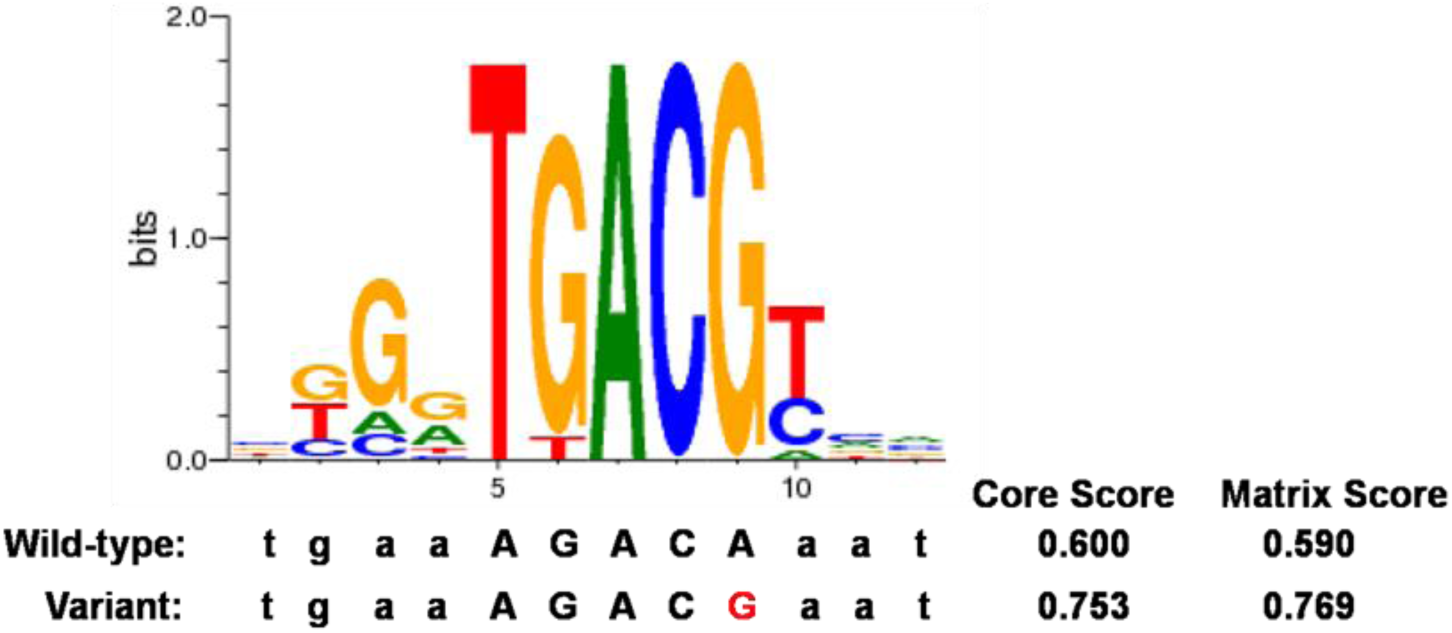
Pictorial representation of matrix for CREB binding motif predicted computationally using P-Match. CREB is predicted to bind to -181G allele with higher affinity.

**Figure S3.**
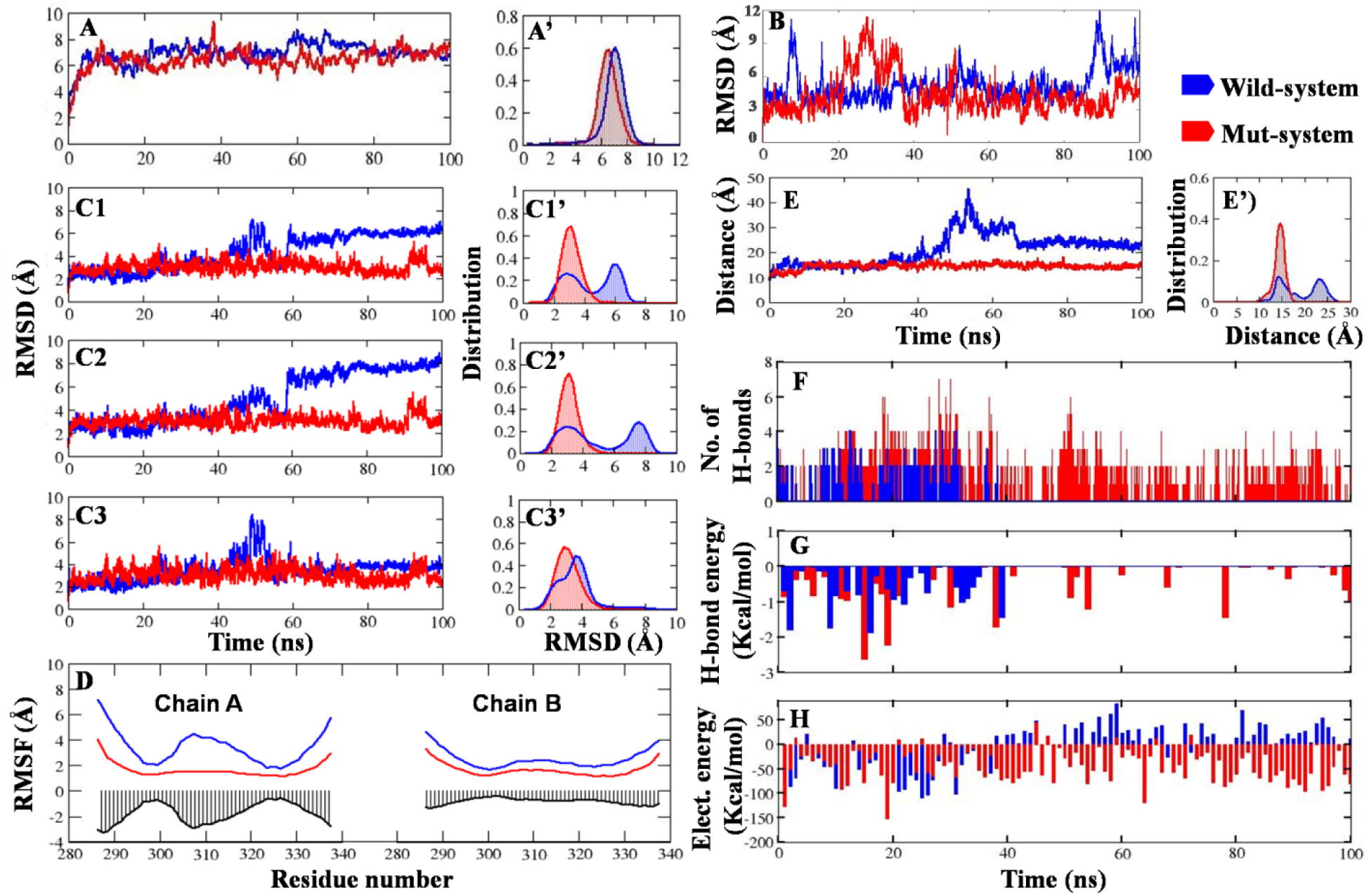
Qualitative and quantitative analysis of wild-type and mut systems: **(A and A’)** RMSD of *MMP7*-promoter DNA (region ±10 bp from -181 bp). **(B)** RMSD of CREB1(monomer)-DNA complex during MD simulations. The CREB1(monomer)-DNA-wild-type system showed an average RMSD of 5.0 Å whereas the CREB1(monomer)-DNA-mut showed as average RMSD of 4.5 Å.RMSD and RMSD distribution profiles from molecular dynamics simulations for CREB1(dimer)-*MMP7*-promoter complexes: C1 and C1’ represent the whole system; C2 and C2’ represent chain A; C3 and C3’ represent chain B. **(D)** RMSF profiles of CREB1(dimer)-DNA complexes. **(E and E’)** The distance vs. time evolution and distance vs. distribution profiles between CREB1 chain A and the polymorphic nucleotide (residue number 23) in chain C in CREB1(dimer)-DNA complex through 100ns MD simulation. **(F, G and H)** Hydrogen bonds occupancy, hydrogen bonds energy and electrostatics energy profiles for nucleotide number 23 in chain C and chain A in CREB1(dimer)-DNA complex.

**Figure S4.**
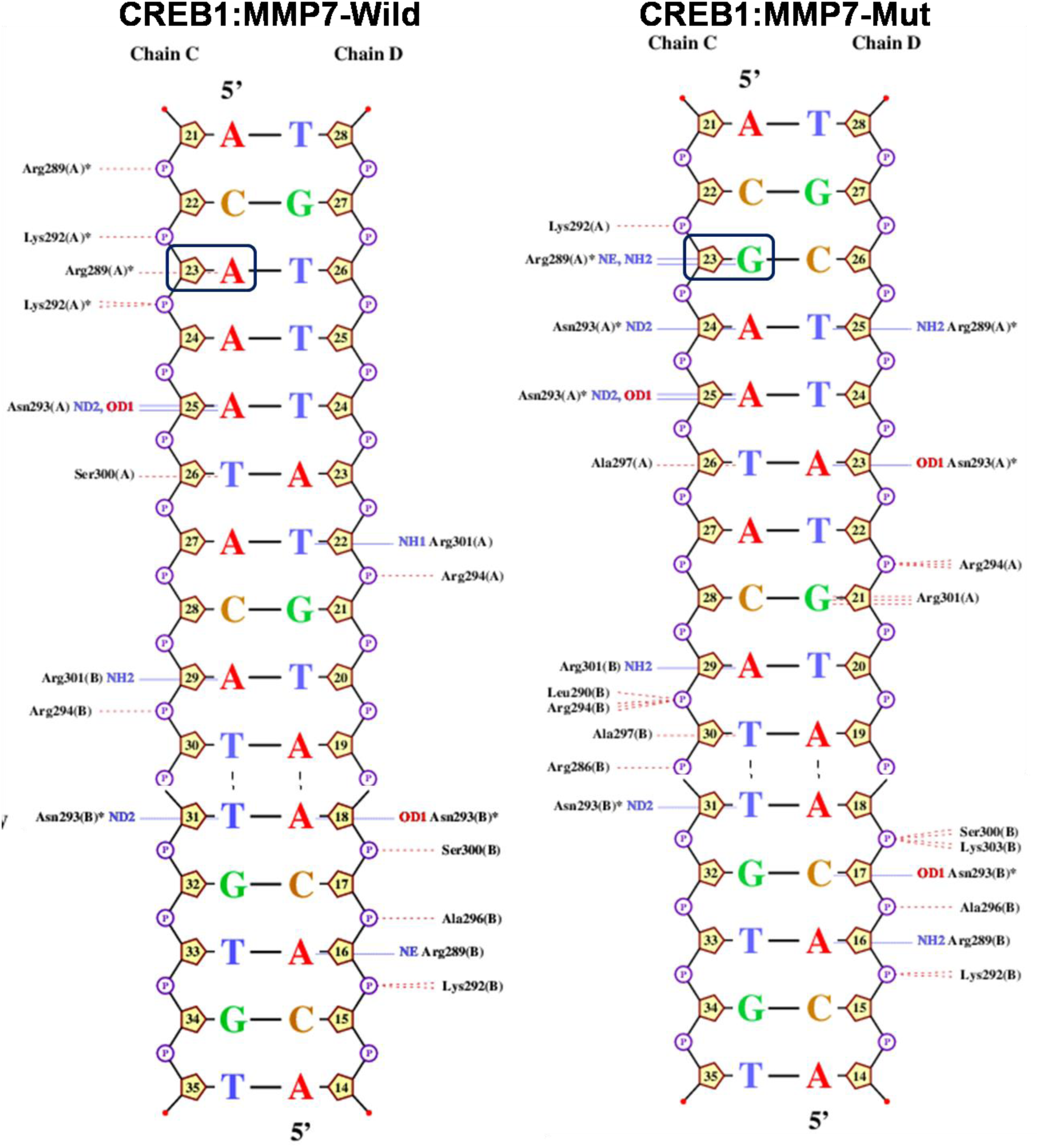
2D-interaction map of protein-DNA complexes of *MMP7*-wild-type:CREB1 and *MMP7*-mutant:CREB1. Hydrogen bonds and van der Waals interactions are represented with solid blue and dotted red lines, respectively. The cut-off distances for hydrogen bonds and van der waals interaction are 3.50 Å and 3.35 Å, respectively. This was generated using NucPlot.

**Figure S5.**
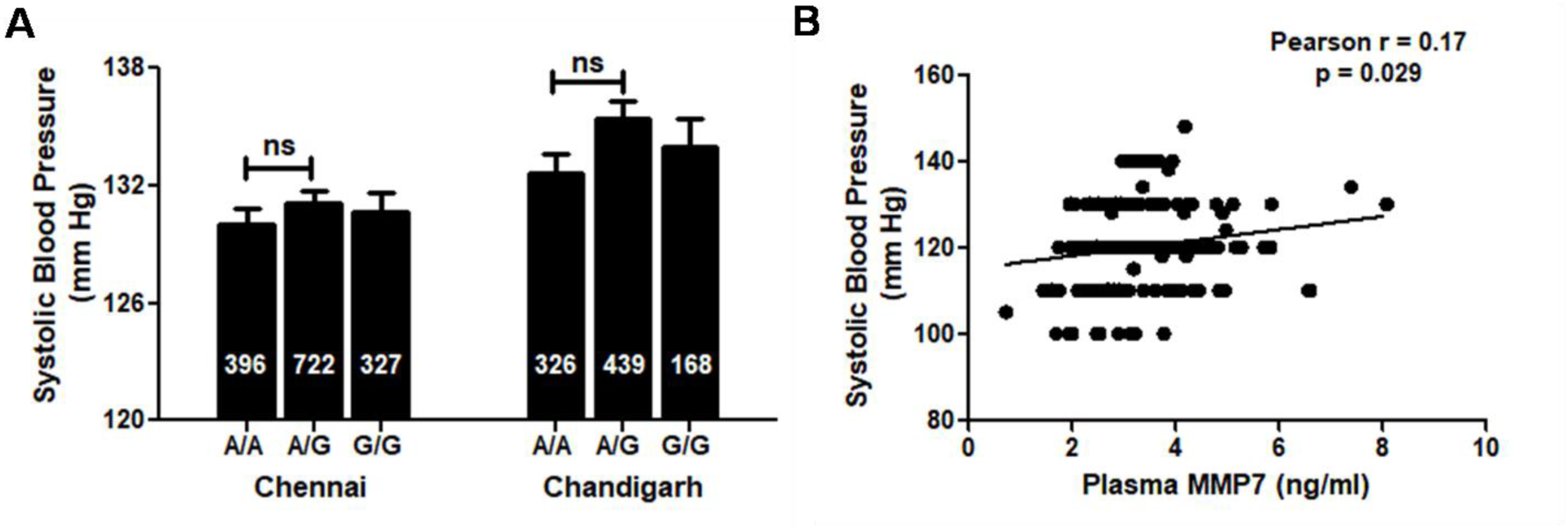
*MMP7*-181 promoter polymorphism and systolic blood pressure. **(A)** *MMP7*-181A/G individuals display a trend towards higher systolic blood pressure in Chennai and Chandigarh populations. **(B)** Correlation of systolic blood pressure with MMP7 levels. MMP7 levels showed significant positive correlation with systolic blood pressure. Pearson r and p values for the correlations are indicated.

